# Pigmentation and retinal pigment epithelium thickness: a study of the phenotypic and genotypic relationships between ocular and extraocular pigmented tissues

**DOI:** 10.1101/2024.12.04.626809

**Authors:** Thomas Julian, Tomas Fitzgerald, UK Biobank Eye and Vision Consortium, Ewan Birney, Panagiotis I. Sergouniotis

## Abstract

The retinal pigment epithelium (RPE) is a specialised monolayer of pigmented epithelial cells in the outer retina. The extent to which RPE pigmentation is related to that of other tissues remains unclear. We utilised RPE thickness measured using optical coherence tomography (OCT) imaging as an indicator of RPE melanin content. UK Biobank data was used to assess the relationships between RPE thickness and fundus pigmentation, hair colour, skin colour, and ability to tan. We performed a genome-wide association study (GWAS) to identify genetic loci associated with RPE thickness. We explored the genetic correlation between RPE thickness and pigmentation-related traits. We found that RPE thickness was not phenotypically or globally genetically correlated with hair colour, skin colour or ability to tan. Whilst RPE thickness was phenotypically correlated with fundus pigmentation, there was not significant global genetic correlation. Despite this, variants in key pigmentation loci including *TYR*, and *OCA2*-*HERC2* were significant in our GWAS of RPE thickness. We identified four genetic regions in which RPE thickness is locally genetically correlated with other pigmentation-related traits, all of which contain protein-coding genes that are central to melanogenesis and melanosome transport. Our study highlights shared and divergent features between RPE thickness and other pigmented traits.

**Significance:** The findings outlined in this study support the assertion that pigmentation is related to RPE thickness and show that whilst RPE thickness’s genetic make-up differs from that of other pigmented traits, there is overlap in genes that have critical roles in pigment production.

**Research Highlights:** Here, we explored the relationships between retinal pigment epithelium (RPE) thickness and pigmented traits within and outside of the eye. We found that: (i) well-known pigmentation genes influence RPE thickness; (ii) the thickness of RPE differs according to skin colour; and (iii) RPE thickness is genetically related to other pigmented traits through genes that influence pigmentation.

## Introduction

The retinal pigment epithelium (RPE) is a single layer of post-mitotic cells located between the photoreceptors and choroid (George *et al*., 2021). It serves multiple functions that are critical to preserving normal retinal physiology including phagocytosis of shed photoreceptor outer segments; transport of nutrients, ions and fluids; maintenance of the visual cycle; and protection from photooxidation by scavenging free radicals and reactive oxygen species (Seagle *et al*., 2005). The latter of these properties is to a large extent dependent upon the pigmented nature of the RPE.

The biology of melanin pigmentation in the RPE differs to that of other pigmented tissues. Unlike the skin and the uveal tract (iris and choroid), RPE melanin is not produced by melanocytes; instead it is synthesised by melanosomes within the RPE itself (Istrate *et al*., 2020). Additionally, the RPE differs from other pigmented cells with regards to its embryological origins. Whilst the pigmented cells of the uvea, the epidermis and the inner ear originate in the neural crest, the RPE is derived from the neural ectoderm (Silver et al., 2013). The initiation of RPE melanin synthesis occurs before other tissues, with pigmentation evident at around the 4th week of gestation (Schneider & Green, 1998). It has been highlighted that melanin synthesis in the RPE is thought to be largely limited to prenatal and perinatal life (Kauffman & Han, 2024), which is at odds with other pigmented tissues, which tend to synthesise melanin throughout life (Benito-Martínez *et al*., 2020).

Optical coherence tomography (OCT) is a method for high-resolution (∼1–3Dµm), cross sectional imaging of tissues (Bouma *et al*., 2022). In OCT, interferometry is used to analyse light reflected from biological structures to construct two- and three-dimensional images of tissues. Because OCT relies on the reflectance of light, the optical properties of the tissues being imaged influence the appearance of the constructed image. Illustratively, in the context of retinal imaging, RPE melanin pigment is known to be responsible for a hyper-reflective OCT band and to cause light scattering effects which impair the resolution of adjacent structures such as Bruch’s membrane and photoreceptor outer segment tips (Meleppat et al., 2019; Wilk et al., 2017). Notably, in zebrafish it has been demonstrated that RPE melanin content influences not only the extent of reflectance of bands on OCT imaging, but also RPE thickness (Wilk *et al*., 2017). Given that RPE is a monolayer, the thickness of the band corresponding to this structure on OCT ought to be influenced by (a) the true height of RPE cells, and (b) the impact of RPE properties (such as melanin content) on the reflectance of light. On an individual basis, the cellular morphology of RPE cells varies according to retinal eccentricity, with taller, thinner cells in the central retina; and wider, flatter cells in the periphery (Boulton & Dayhaw-Barker, 2001). On a population level, the OCT-measured thickness of the RPE has been shown to be influenced by age, ethnic group and refractive error (Ko *et al*., 2017; Shao *et al*., 2022). The causal pathways underlying the relationship between ethnicity and RPE thickness has not been fully explored, but could be underpinned by tissue pigmentation.

Collectively, existing evidence suggests that pigmentation has a substantial impact on the OCT-measured thickness of the RPE, and therefore its thickness is likely to correlate with pigmentation of the tissue. In this study, we seek to explore the association between OCT-measured RPE thickness and a variety of pigmentation-related traits. By exploring phenotypic and genotypic relationships between a RPE thickness and pigmentation, we seek to identify shared and divergent biological features of the RPE relative to other pigmented tissues. Finally, utilising the genetic data generated in this study, we explore causal determinants of RPE thickness using Mendelian randomisation.

## Methods

### Ethics statement

The UK Biobank (UKB) study was conducted with the approval of the North-West Research Ethics Committee (ref 06/MRE08/65), in accordance with the principles of the Declaration of Helsinki, all participants gave written informed consent and were free to withdraw at any time (Sudlow *et al*., 2015). This project used data from the UKB study under approved project numbers 53144 and 49978.

### Cohort characteristics

UKB is a prospective population-based study of 502,355 participants. Baseline examinations were carried out at 22 study assessment centres between January 2006 and October 2010. Study participants have undergone a detailed assessment of demographic, lifestyle and clinical measures; provided DNA samples via blood tests; and provided a range of physical measures. A substantial subset of volunteers underwent an ophthalmic assessment (23%, ND=D117,279). Of these subjects, 67,664 individuals underwent spectral domain OCT between 2006-2010 (‘instance 0’) and an additional 17,090 participants were imaged for the first time during the first repeat assessment between 2012-13 (“Instance 1”) (Littlejohns *et al*., 2020).

### OCT phenotypes

OCT images in UKB have previously been segmented using the Topcon Advanced Boundary Segmentation algorithm (Ko *et al*., 2017). We selected our primary RPE trait for analysis on the basis of population size and image quality. Accordingly, the metric studied in our main analysis was left eye overall average RPE thickness (UKB data field 27822). This is the derived RPE thickness value with the largest sample size (80,305 participants before quality control). Furthermore, given that the left eye was consistently imaged second, it is likely that the left eye image would, on average, be of higher quality as both the subject and technician would be more prepared. This assertion is supported by a marginally higher median image quality metric in left eye images than right eye images (68 vs 67). Additional RPE traits considered in this study were the RPE thickness across retinal subfields, inclusive of the left and right eye central subfields, inner subfields and outer subfields. The central subfield was defined as per the Early Treatment Diabetic Retinopathy Study (ETDRS) grid (“Grading Diabetic Retinopathy from Stereoscopic Color Fundus Photographs—An Extension of the Modified Airlie House Classification: ETDRS Report Number 10,” 1991). The inner retinal subfield was defined as the mean value across the inner nasal, temporal, superior and inferior subfields; whilst the outer retinal subfield was the mean of the outer nasal, temporal, superior and inferior subfields. As illustrated in Supplementary Figures S1-9, the RPE values for these phenotypes were positively skewed.

### Retinal pigmentation score (RPS)

The optical density of melanin is substantially greater in the choroid than the RPE, and therefore it is believed that the choroid is the main contributor of visible pigmentation of the fundus (Weiter *et al*., 1986). In order to quantify the extent of pigmentation of the posterior pole, we utilised the retinal pigmentation score (RPS) model developed by Rajesh et al (Rajesh *et al*., 2023). It is likely that this score is principally a measure of choroidal pigmentation. The methodology employed in this model is documented in the author’s GitHub page (https://github.com/uw-biomedical-ml/retinal-pigmentation-score) (Rajesh *et al*., 2023). Using the developers’ recommendations, we calculated the RPS in 32,224 UKB subjects who had sufficiently high quality fundus photographs according to the “Good” quality filter in Automorph (Rajesh et al., 2023; Zhou et al., 2022). The RPS was skewed (Supplementary Figure 10), and so values were rank-based inverse normalised prior to GWAS but raw values were used in conventional epidemiological analyses. 18,471 subjects with an RPS were available for the genetic analysis following quality control detailed below.

### OCT quality control

In keeping with previous studies, we excluded all images with image quality scores less than 40 and images representing the poorest 10% as designated by the inner limiting membrane (ILM) indicator. We also excluded any image with a RPE layer thickness greater than 2.5 standard deviations from the mean (Zekavat *et al*., 2022). This ensured that mostly high-quality images remained in our analysis, reducing the risk of including OCT scans with segmentation errors.

### Phenotypic statistical analysis

Non-genetic statistical tests were conducted for subjects with complete data with respect to left eye overall average RPE thickness, RPS, hair colour (data field 1747), skin colour (data field 1717), and tanning ability (data field 1727). The cohort was not filtered according to ethnicity/ancestry for this analysis. For this phenotypic analysis, the hair colour, skin colour and tanning ability were treated as ordinal numerical variables using the default UKB coding systems described in the UKB showcase directory (*UK Biobank*, n.d.), whilst RPE thickness and RPS were treated as continuous numerical variables. The Kruskal Wallis rank sum test followed by pairwise comparisons using Wilcoxon sum rank test was then used to look for differences between the studied phenotypes. Phenotypic correlations were calculated using Kendall’s Tau.

### Genetic association studies

Genome wide association study (GWAS) was performed using an additive linear model implemented in REGENIE v3.1.1. Given the normality assumption of GWAS, we performed rank-based inverse normal transformation of the derived OCT values prior to analysis (Mbatchou *et al*., 2021). Subjects who were related to the third degree or more; outliers for heterozygosity or missing rate; and those with sex chromosome aneuploidy were excluded. The following quality-control filters were applied on the imputed genotype data (UK Biobank data-field 22828) using PLINK (Purcell *et al*., 2007): a minor allele frequency (MAF) >5%; Hardy–Weinberg equilibrium test not exceeding 1 × 10^−15^; a genotyping rate above 99%; not present in a low-complexity region, a region of long-range linkage disequilibrium or a sex chromosome. For our RPS GWAS, the same protocol was followed, except variants with a minor allele count <20 were additionally excluded and a genotype rate above 90% was utilised. Correction for the following covariates was undertaken: age at recruitment (data-field 21022), sex (data-field 31), height (data-field 50), weight (data-field 21002), refractive error (calculated as spherical error + 0.5 × cylindrical error; data-fields 5085 and 5086) and genetic principal components 1 to 20 (data-field 22009). A conventional genome-wide significance p value of p<5E-08 was selected for the identification of genetic variants associated with our primary outcome. Following both genetic and image quality control, our sample sizes for genetic studies were as follows: left eye overall RPE thickness 29,550; right eye overall RPE thickness 29,309; left eye mean outer RPE thickness 8,840; right eye mean outer RPE thickness 8,505; left eye inner RPE thickness 9,036; right eye inner RPE thickness 8,657; left eye central RPE thickness 9,331; right eye central RPE thickness 8,983; replication left eye overall RPE thickness 6,727; and RPS 18,471.

To refine the obtained association signals, further analyses were performed using the GCTA-COJO tool (https://yanglab.westlake.edu.cn/software/gcta/#COJO) (Purcell *et al*., 2007; Yang *et al*., 2011). These analyses were conducted utilising linkage disequilibrium estimates from a reference sample and summary statistics from each phenotype described previously (Currant *et al*., 2021). Genetic variants in loci that were on different chromosomes or more than 10 Mb distant from each other were assumed to be uncorrelated.

### Mendelian randomisation

Causal associations between 10,310 exposures and our primary metric (left eye overall average RPE thickness) were assessed using univariable two-sample Mendelian randomisation. The methodology of the utilised phenome-wide Mendelian randomisation pipeline has been described in detail elsewhere (Julian, Cooper-Knock, *et al*., 2023; Julian, Girach, *et al*., 2023). The current analysis differs only in the selection of exposures from the IEU open GWAS database and it is noted that exposures in the prot-a, prot-b, prot-c, met-a, met-c, ieu-b, and ebi-a datasets were included. Descriptions of these datasets are available via the IEU OpenGWAS project (*IEU OpenGWAS project*, n.d.). “Non-European” ancestry GWAS were excluded, because two sample Mendelian randomisation assumes that the data are derived from the same broad ancestral population (to avoid confounding by differing population structure). We defined causal traits to be those which were: significant after false discovery rate (FDR) correction in the multiplicative random effects inverse variance weighted (MRE IVW) analysis; nominally significant in the weighted median (Bowden *et al*., 2016), nominally significant in the weighted mode (Bowden et al., 2016; Hartwig et al., 2017) and nominally significant throughout a leave one out analysis using the MRE IVW. Population overlap biases univariable two-sample MR in the direction of causation. Accordingly, the populations contributing toward traits determined to be causal were manually inspected to ensure that there was not considerable overlap (i.e. exposure the cohort was largely constituent of UKB subjects). The results presented in the main manuscript have no sample overlap while the results presented in the supplementary content (*i.e.* 10,310 Mendelian randomisation findings) have not been manually inspected for overlap.

### Global genetic correlation analysis

The global genetic correlation between left eye overall average RPE thickness and pigmentation-related traits including iris colour, RPS, hair colour, skin colour and ability to tan was assessed using linkage disequilibrium (LD) score regression (Bulik-Sullivan *et al*., 2015). The GWAS statistics for the studied pigmentation-related traits were either generated as part of this study or derived from previously published studies (Backman et al., 2021; Loh et al., 2018; Simcoe et al., 2021). This analysis was consistent with the methodology outlined by Bulik-Sullivan *et al*., in which genetic variants were filtered according to: presence in HapMap3; MAF > 0.01; INFO score >0.9; removal of strand-ambiguous variants and duplicated variants. LD scores were derived from the European subset of the 1000 Genomes Project dataset (1000 Genomes Project Consortium *et al*., 2015). The intercept was not constrained in this analysis. LD score regression was also utilised to determine the heritability of the left eye overall average RPE thickness trait.

### Local genetic correlation analysis

The local genetic correlation between left eye overall average RPE thickness and the pigmentation-related traits described above was calculated using Local genetic correlation analysis using Local Analysis of [co]Variant Association (LAVA) (Bulik-Sullivan *et al*., 2015; Werme *et al*., 2022). The sample overlap was estimated using the intercepts derived from LD score regression as described above. The European subset of the 1000 Genomes Project dataset was utilised for estimation of LD. Loci of interest were defined in the same manner described by Werme et al. (Werme *et al*., 2022), and, as a result, the pairwise local correlation between traits was assessed at 2,495 genomic locations. The detection of stable and interpretable local genetic correlation requires sufficient local genetic signal. As such, the local heritability at each locus for each trait was calculated, and bivariate analysis was only carried out at a given locus where two traits each exhibited multiple testing corrected levels of significant heritability (i.e., p<0.05/2,495). In the bivariate analysis, our primary outcome was the local genetic correlation between RPE and pigmentation-related traits. The threshold for identification of significant relationships was Bonferroni corrected (0.05/1,060 bivariate tests).

### Software

GWAS was performed using REGENIE (version 3.1.1). COJO was performed using the GCTA COJO tool (version 1.94.1). PLINK (version 1.9) was used for variant filtering. MR was performed using the TwoSampleMR package (version 0.5.7) in R (version 4.2.2). LAVA was performed using the LAVA package (version 0.1.0) in R. LD score regression was performed using the LDSC package (version 1.0.1) in Anaconda, Python (version 2.5.0). Regression was conducted using the stats package (version 4.2.2) in R. Scatter and correlation plots were made using the ggplot2 package in R. Histograms were constructed using ggplot in R. Q-Q plots and manhattan plots were constructed using the qqman package in R. LAVA chord plots were constructed using the circlize package (version 0.4.15) in R. Network plots were made using igraph (version 2.1.1). The packages required for calculation of the RPS are detailed on the model GitHub page (Rajesh *et al*., 2023).

## Results

### Phenotypic statistical analysis

Overall, 26,590 subjects were included in the phenotypic analysis. In terms of self-identified broad ethnic groups, 24,990 selected White, 489 Black, 528 Asian, 220 Mixed, 70 Chinese and 293 other ethnicities. The mean overall left eye RPE thickness was found to be 24.67µm, with an interquartile range of 3.65µm. Although there was a significant (p=1.44e-06) negative correlation between RPE thickness and RPS, the strength of correlation was very weak (Kendall’s Tau coefficient -0.02) and therefore of little practical significance, as can be seen in Supplementary Figure S11. Although there was a nominally significant weak negative correlation between RPE thickness and tanning ability (p=0.03), the significance did not exceed multiple testing correction (Figure 1). There was no significant phenotypic correlation between RPE thickness and hair colour or skin colour. The coefficients of Kendall’s Tau are presented in Supplementary Table 1.

**Figure 1:**
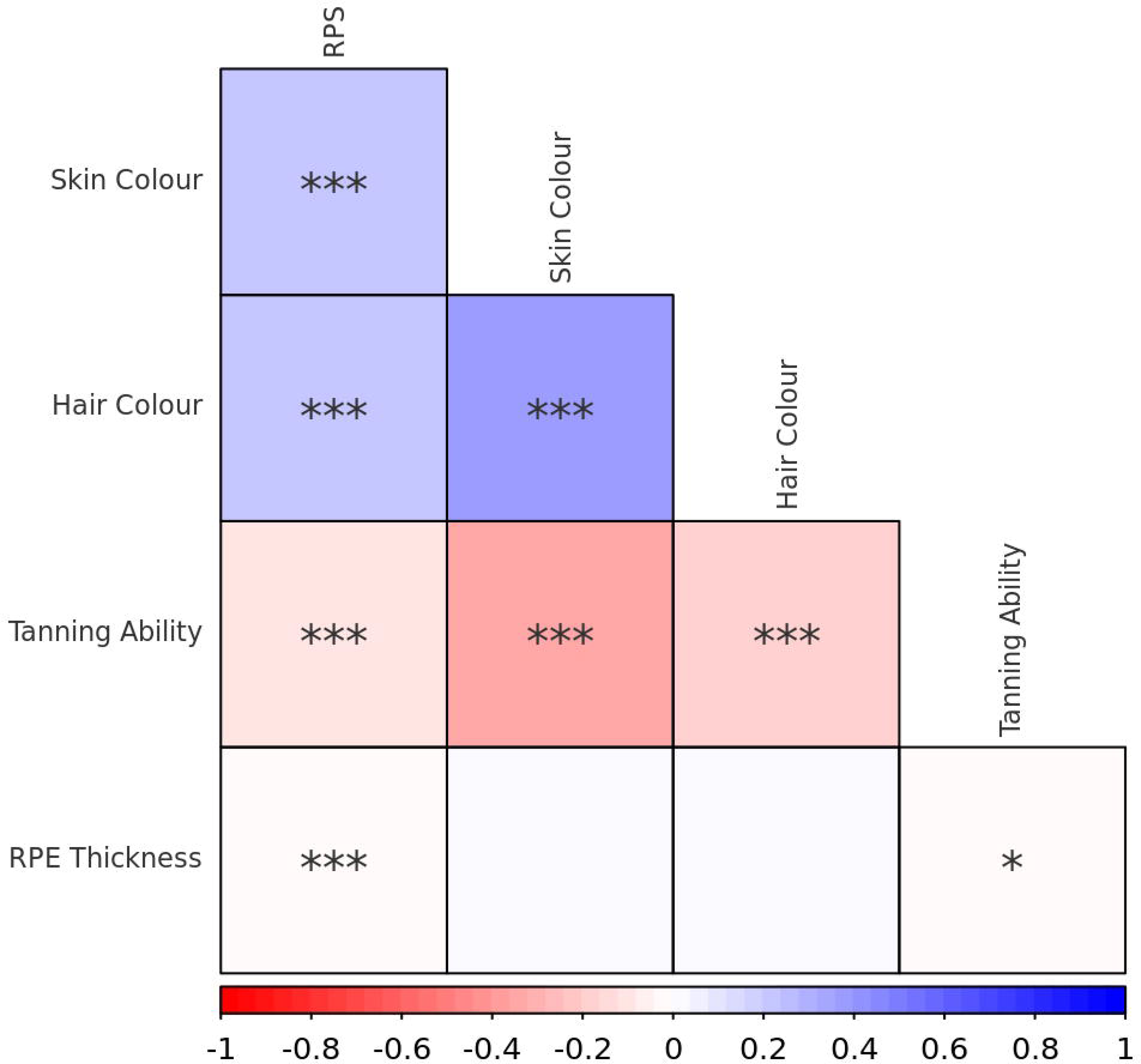
A phenotypic correlation plot of pigmentation-related traits available in UK Biobank. Red colour indicates a negative correlation, whilst blue colour indicates a positive correlation. RPE, retinal pigment epithelium; RPS, retinal pigmentation score. * = p<0.05, ** = 0<0.01, *** = p<0.001.

The “RPS” is a deep learning score which principally reflects the pigmentation of the choroid, which is the main contributor to fundus pigmentation. The distribution of RPE thickness and RPS varied significantly according to skin colour (Kruskal-Wallis rank sum test p-value <2.2e-16 in both cases). For RPE thickness, the Wilcoxon rank sum test indicates that the signal is driven by significant differences between those who reported having brown or black skin, and those who reported having fair, light olive or dark olive skin. For RPS, there were significant differences between all skin colour pairwise comparisons (Supplementary Table S2). These results are visually supported by the box and whisker plots presented in Figures 2 & 3, and Supplementary Figures S12-13. The distribution of RPE thickness also varies according to the self-reported ethnic group, as can be seen in Supplementary Figures 14-15.

**Figure 2:**
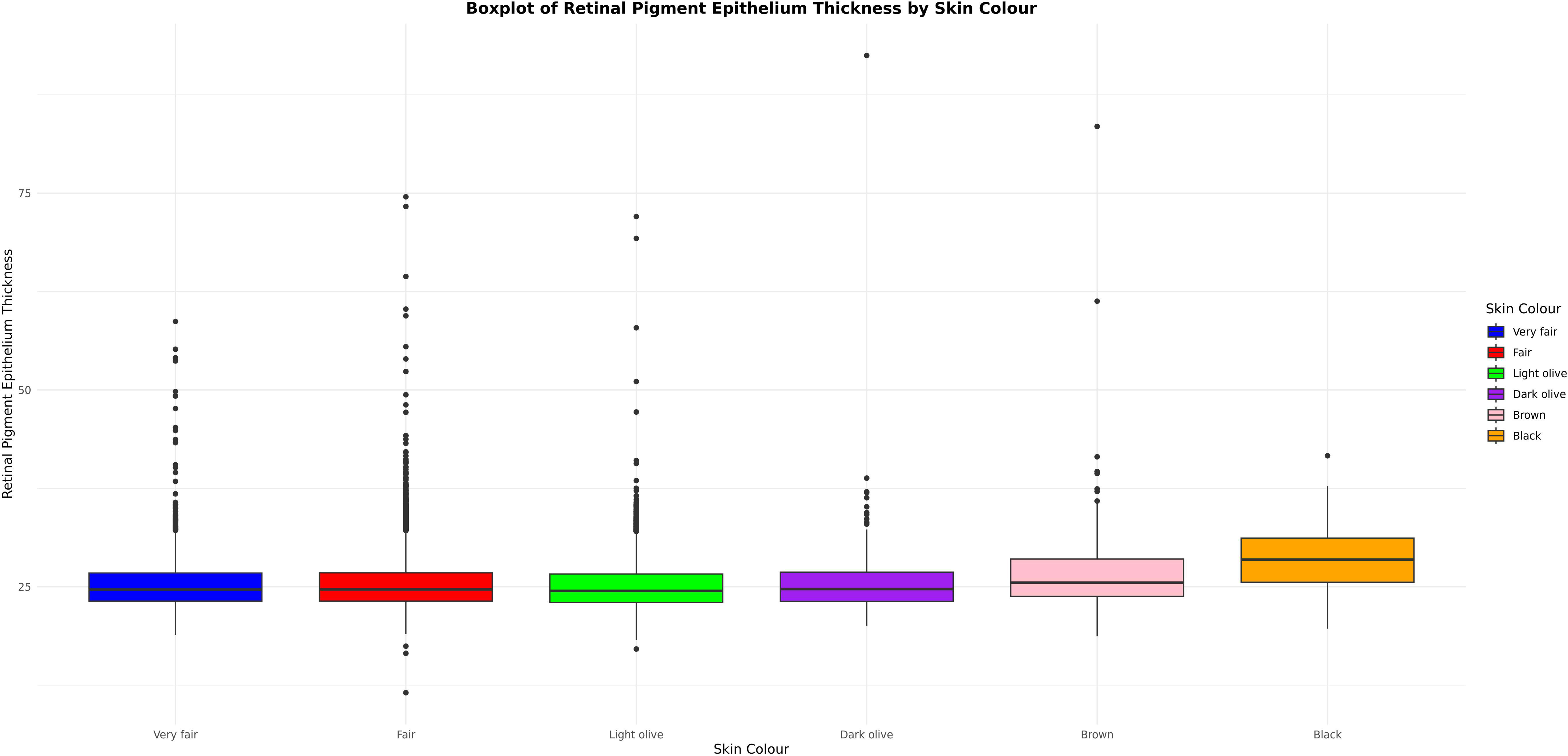
A box and whisker plot illustrating the distribution of left eye overall average RPE thickness according to self-reported skin pigmentation.

**Figure 3:**
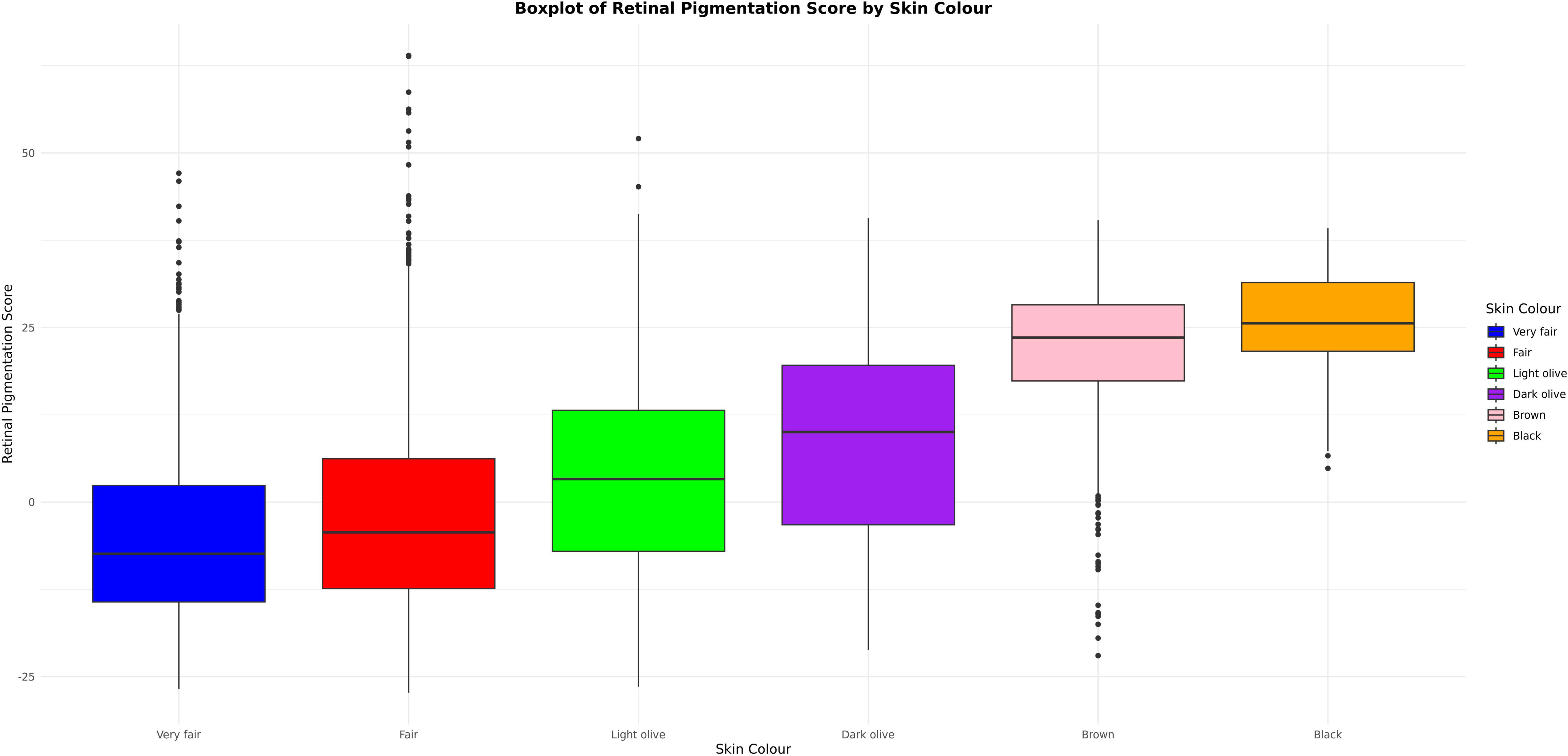
A box and whisker plot illustrating the distribution of RPS according to self-reported skin pigmentation.

## Genetic analyses

### Genetic association studies

In our primary analysis of left eye overall average RPE thickness, GWAS results were obtained for 1,691,495 variants for 29,550 individuals. LD score regression indicated that the h^2^ of left eye overall RPE thickness was 0.21, and that the LD score regression intercept was 0.998. In total, 3,301 genetic variants were found to be associated with left eye overall RPE thickness at a genome-wide significance level (Figure 4). Of the genome-wide significant variants, GCTA-COJO identified 21 variants that were independently associated with left eye overall average RPE thickness (Table 1). Three of these SNPs are also genome-wide significant (p<5E-08) in every other retinal field in both the left and right eyes (rs1126809 [*TYR*], rs1800407 [*OCA2*] and rs12913832 [*OCA2-HERC2*]), whilst a broader set of 8 further variants were at least nominally significant in every other subfield GWAS (rs377695367 [*GDPD4*], rs62075723 [*NPLOC4-TSPAN10*], rs1410996 [*CFHR3*], rs443134 [*CFH-CFHR3*], rs3770526 [*MREG*], rs2625955 [*RHO*], rs4840499 [*PRSS51*] and rs57819090 [*C8orf74*]). Our replication GWAS for left eye overall average RPE thickness had a relatively small sample size (N=6,727). 3 genetic variants from our primary discovery GCTA-COJO analysis were genome-wide significant in our replication study (rs1126809 [*TYR*], rs12913832 [*OCA2-HERC2*], and rs1410996 [*CFHR3*]). 11 further variants were at least nominally significant (rs61899197 [*SLC15A3*], rs1800407 [*OCA2*], rs62075723 [*NPLOC4*-*TSPAN10*], rs752056016 [*AMPD2*], rs443134 [*CFHR3/CFH*], rs373079331[*NUAK2*], rs3770526 [*MREG*], rs2625955 [*RHO*], 6:1957051_CT_C [*GMDS*], 6:42661637_TTTTC_T [Intronic], rs57819090 [*C8orf74*]. The beta-beta plot provided in Supplementary Figure S16 suggested good agreement between the discovery and replication GWAS in terms of effect size (Pearson coefficient 0.93, p<2.2e-16). The Manhattan plots and QQ plots for all analyses are presented in Supplementary Figures S17-34.

**Figure 4:**
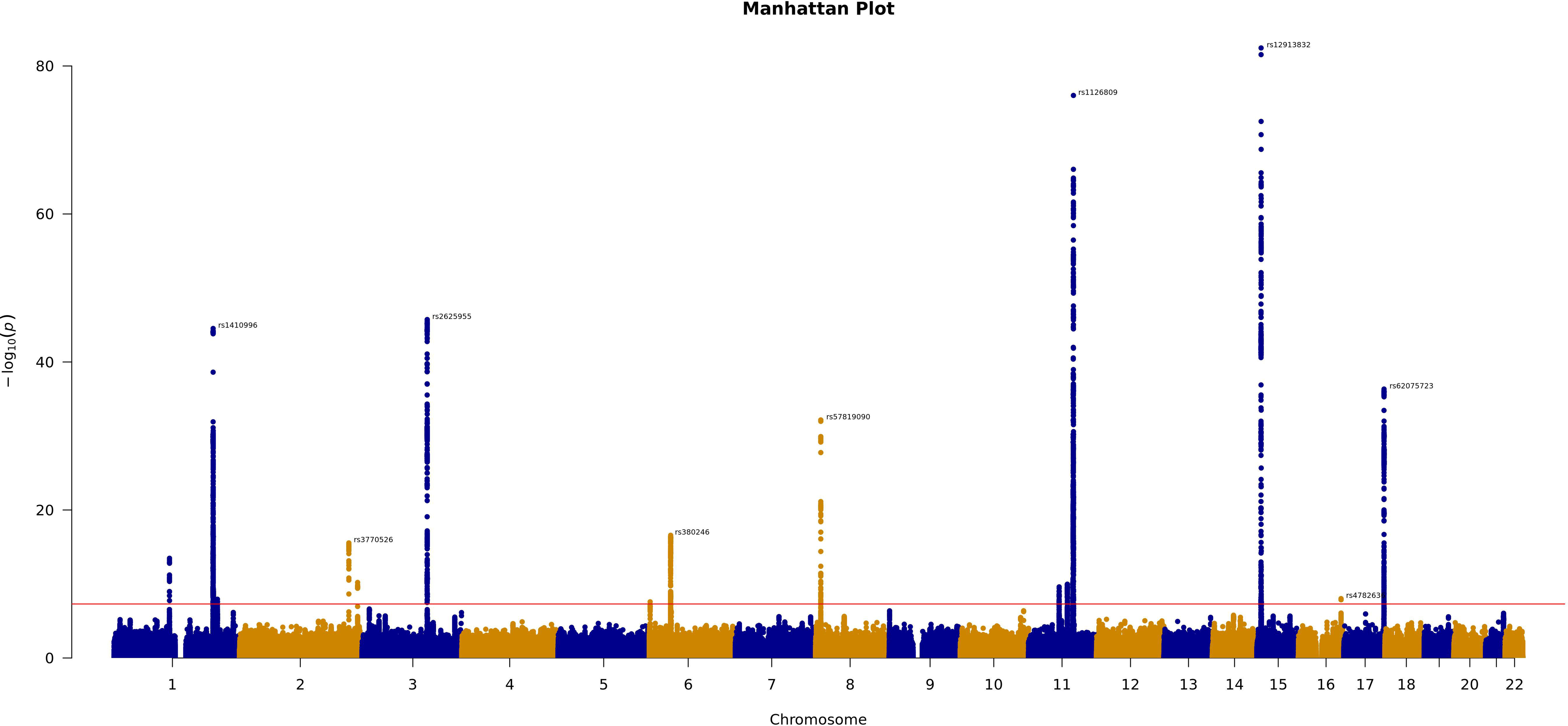
A Manhattan plot, showing the results of a GWAS of left eye overall average RPE thickness, our primary analysis. The red line indicates the threshold for genome-wide significance and key lead genetic variants above this value have been annotated.

**Table 1:** Fine-mapped GWAS results for the primary metric of left eye overall average RPE thickness. Where a variant does not have a rsid, the variant is written in the format chromosome:position_other allele_effect allele.

| Genetic variant [rs ID] | Gene | Effect allele | Joint analysis beta | Joint analysis p-value |
| --- | --- | --- | --- | --- |
| rs61899197 | *SLC15A3* | A | 0.06 | 2.52e-10 |
| rs377695367 | *GDPD4* | A | 0.05 | 1.10e-10 |
| rs1126809 | *TYR* | A | 0.16 | 7.11e-76 |
| rs17566952 | *OCA2* | G | -0.12 | 6.95e-14 |
| rs1800407 | *OCA2* | T | -0.19 | 5.83e-37 |
| rs12913832 | *OCA2-HERC2* | G | 0.13 | 1.35e-39 |
| rs551348273 | *GOLGA8F* | C | 0.08 | 1.90e-12 |
| rs4782636 | *MEAK7* | T | 0.06 | 9.63e-09 |
| rs62075723 | *NPLOC4-TSPAN10* | A | 0.10 | 7.08e-37 |
| rs752056016 | *AMPD2* | A | -0.06 | 3.54e-14 |
| rs1410996 | *CFHR3* | A | 0.09 | 1.59e-23 |
| rs443134 | *CFHR3/CFH* | A | 0.07 | 4.27e-10 |
| rs373079331 | *NUAK2* | CA | -0.07 | 1.17e-08 |
| rs3770526 | *MREG* | T | 0.11 | 3.05e-16 |
| rs55895356 | *SAG* | T | 0.11 | 6.41e-11 |
| rs2625955 | *RHO* | A | 0.15 | 3.75e-46 |
| 6:1957051_CT_C | *GMDS* | C | -0.06 | 2.61e-08 |
| 6:42661637_TTTTC_T | Intronic | T | 0.05 | 6.08e-09 |
| rs380246 | *PRPH2* | T | 0.06 | 2.77e-10 |
| rs4840499 | *PRSS51* | C | 0.06 | 2.97e-11 |
| rs57819090 | *C8orf74* | A | -0.12 | 2.31e-23 |

### Global genetic correlation

LD score regression demonstrated that there are highly significant relationships between hair colour, skin colour, and tanning ability, but not between RPE thickness and other pigmentation-related traits (Figure 5). Whilst there was nominally significant global correlation between RPE thickness and blonde hair colour (p=0.04), this did not pass Bonferroni multiple testing correction. Similarly, RPS and iris colour were not significantly related to other pigmentation-related traits after multiple testing correction. Unfortunately, due to data sharing agreements, we were only able to acquire the Visigen cohort eye colour GWAS summary statistics, and the resulting reduced sample size and missing variants precluded calculation of genetic correlation between iris colour and RPE thickness or RPS. Interestingly, no significant global genetic correlation was identified between RPE thickness and RPS.

**Figure 5:**
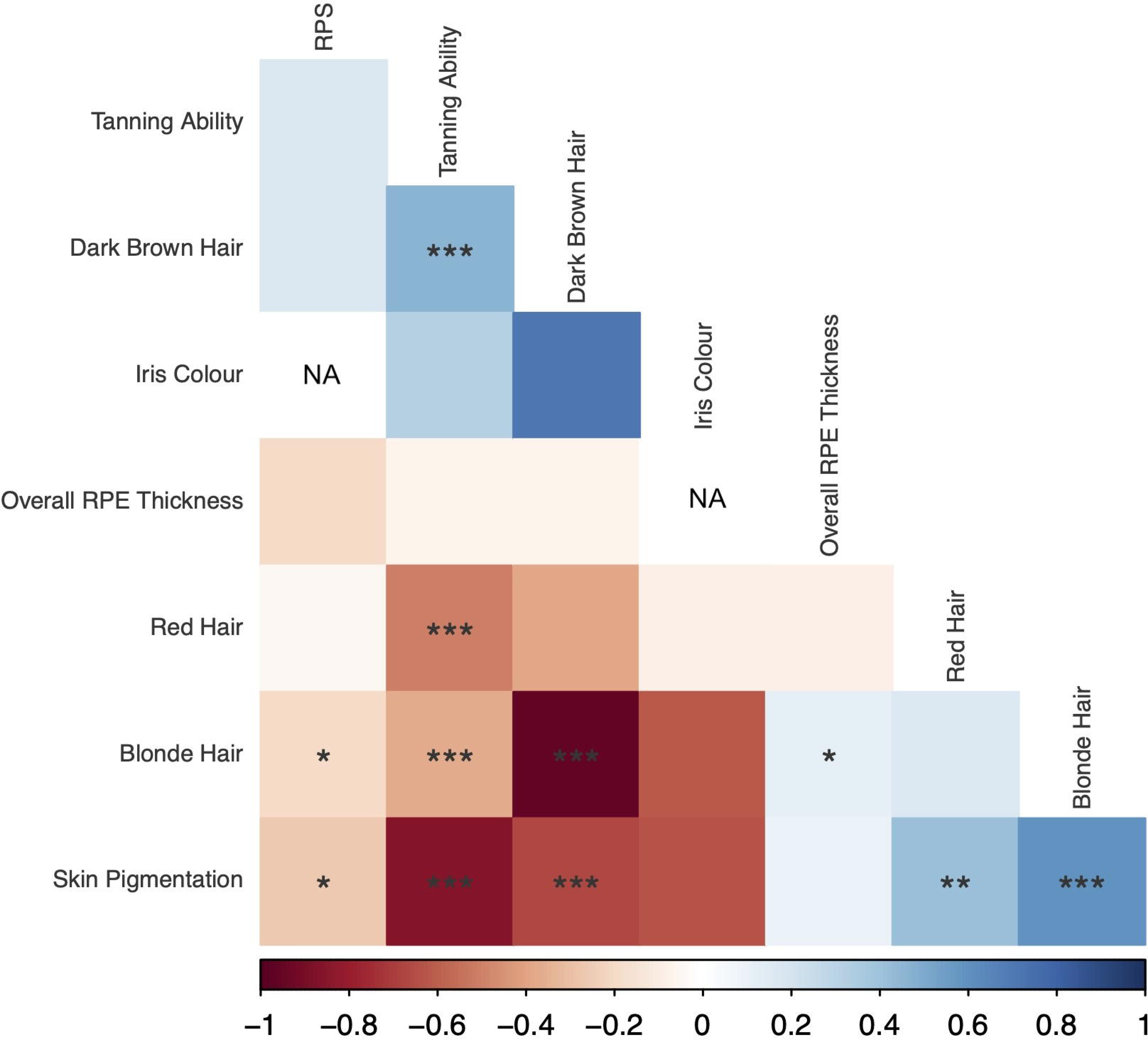
A correlation plot for the global genetic correlation between pigmentation-related traits, assessed using LD score regression. Red colour indicates a negative correlation, whilst blue colour indicates a positive correlation. RPE, retinal pigment epithelium; RPS, retinal pigmentation score. * = p<0.05, ** = p<0.01, *** = p<0.001.

### Local Genetic Correlation

Although there were no significant global genetic relationships between RPE thickness and other ocular or extraocular pigmentation-related traits, there were four regions of highly significant correlation at a local level (Figure 6 and Table 2). Unsurprisingly, these loci contain genes that have been implicated in pigmentation both in biological terms and statistically in previous GWAS (for instance *TYR*, *RAB38*, *OCA2*-*HERC2*, *TSPAN10*, and *HERC2P11*). Network plots are provided to visually illustrate the multiple testing corrected significant genetic relationships in the loci containing *TYR* and *OCA2-HERC2* in Supplementary Figures S35 and S36 respectively. The full results of LAVA local correlation analysis can be viewed in Supplementary Table S3.

**Figure 6:**
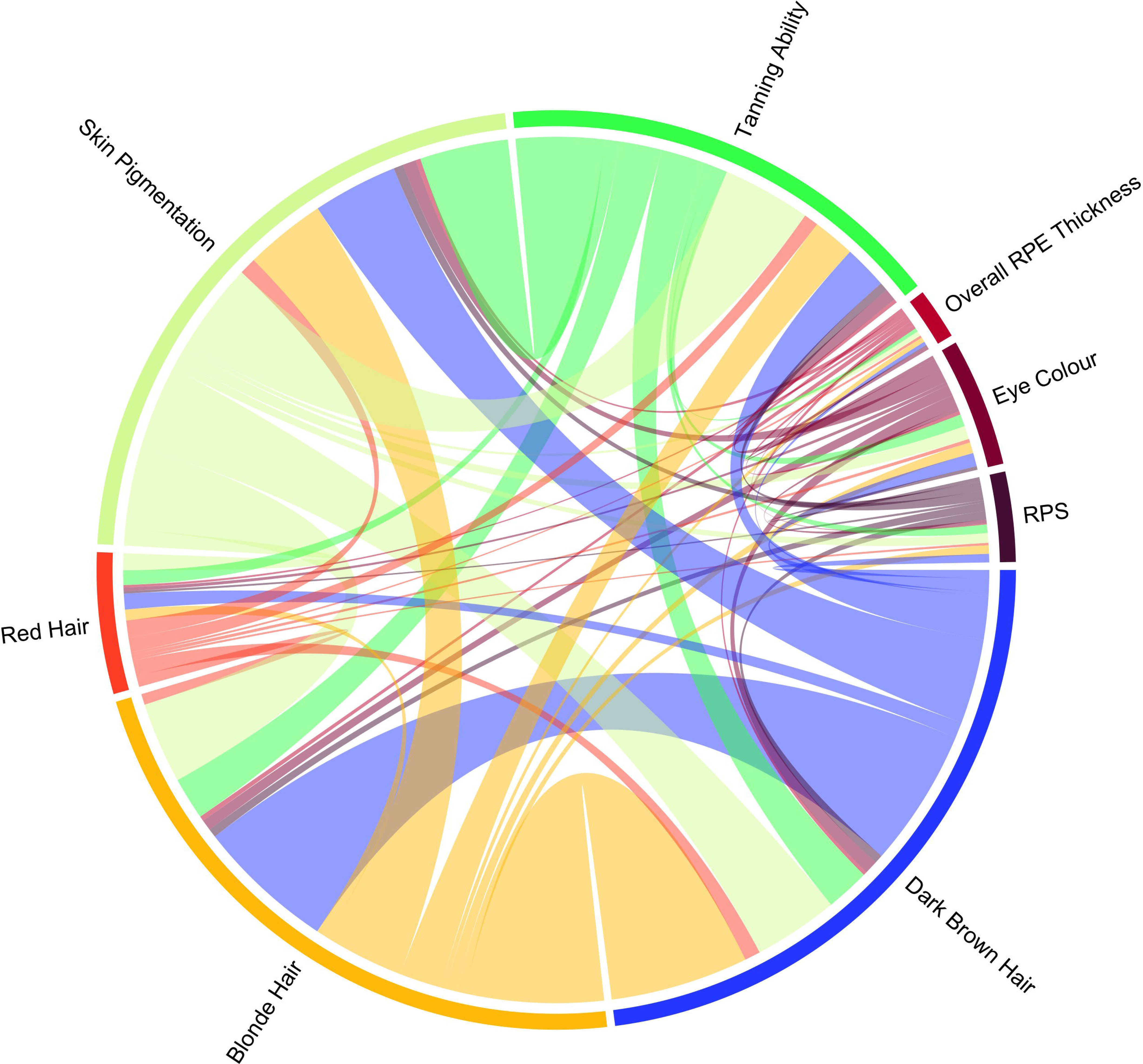
Chord diagram showing the number of multiple testing corrected statistically significant relationships present between key pigmentation-related traits. The coloured connections between traits indicate significant relationships; the wider the connecting line the larger the number of statistically significant local correlations that were identified. Our findings indicate that whilst RPE thickness is genetically correlated with other pigmented tissues in genetic blocks containing pigmentation genes, the overall genetic architecture diverges from that of other tissues. The results suggest whilst there is not global correlation between RPE thickness and other pigmented phenotypes, there is highly significant local correlation in pigment specific regions. We wonder if this may in part be due to the differing embryological origins of the pigmented tissues

**Table 2:**
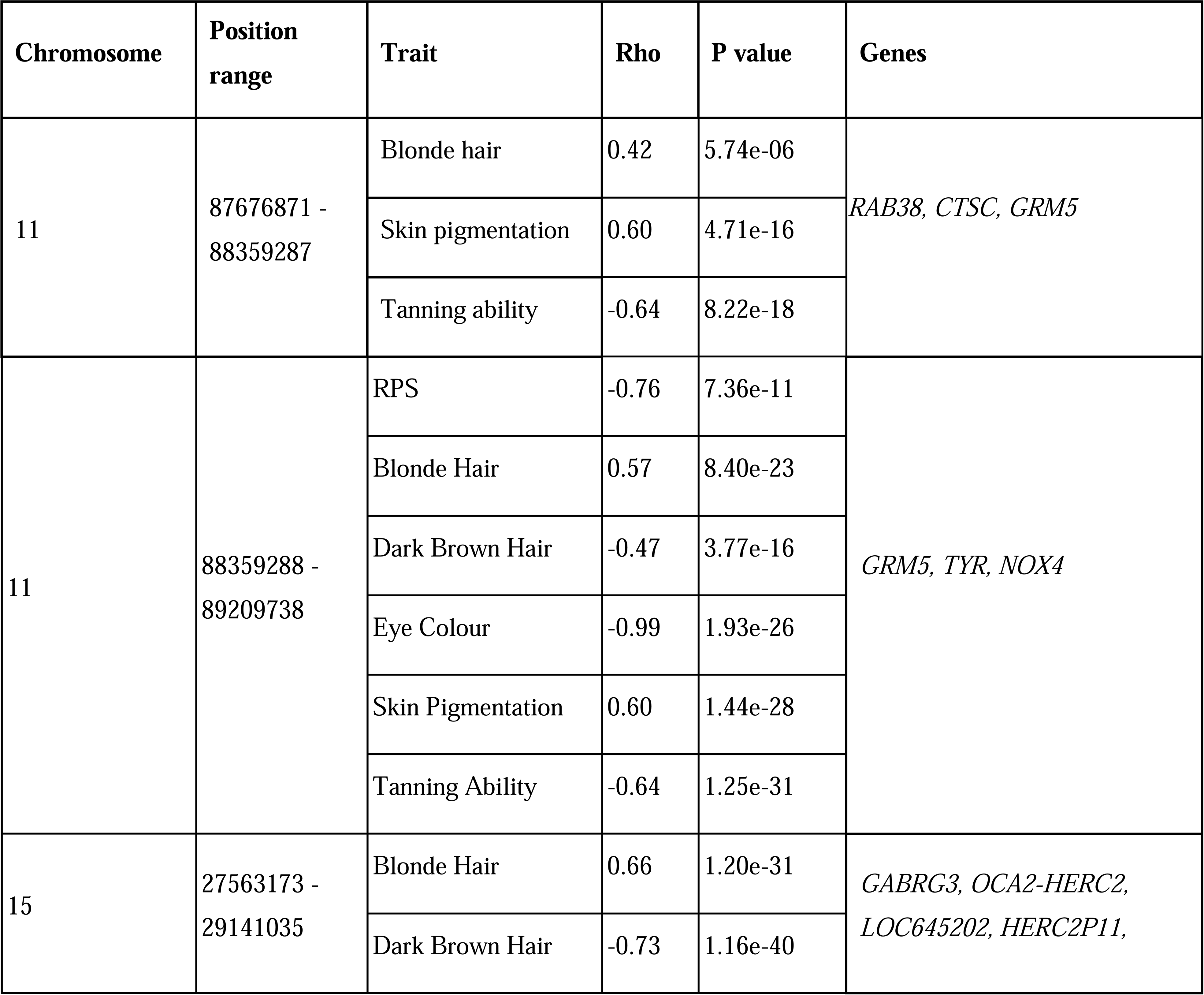
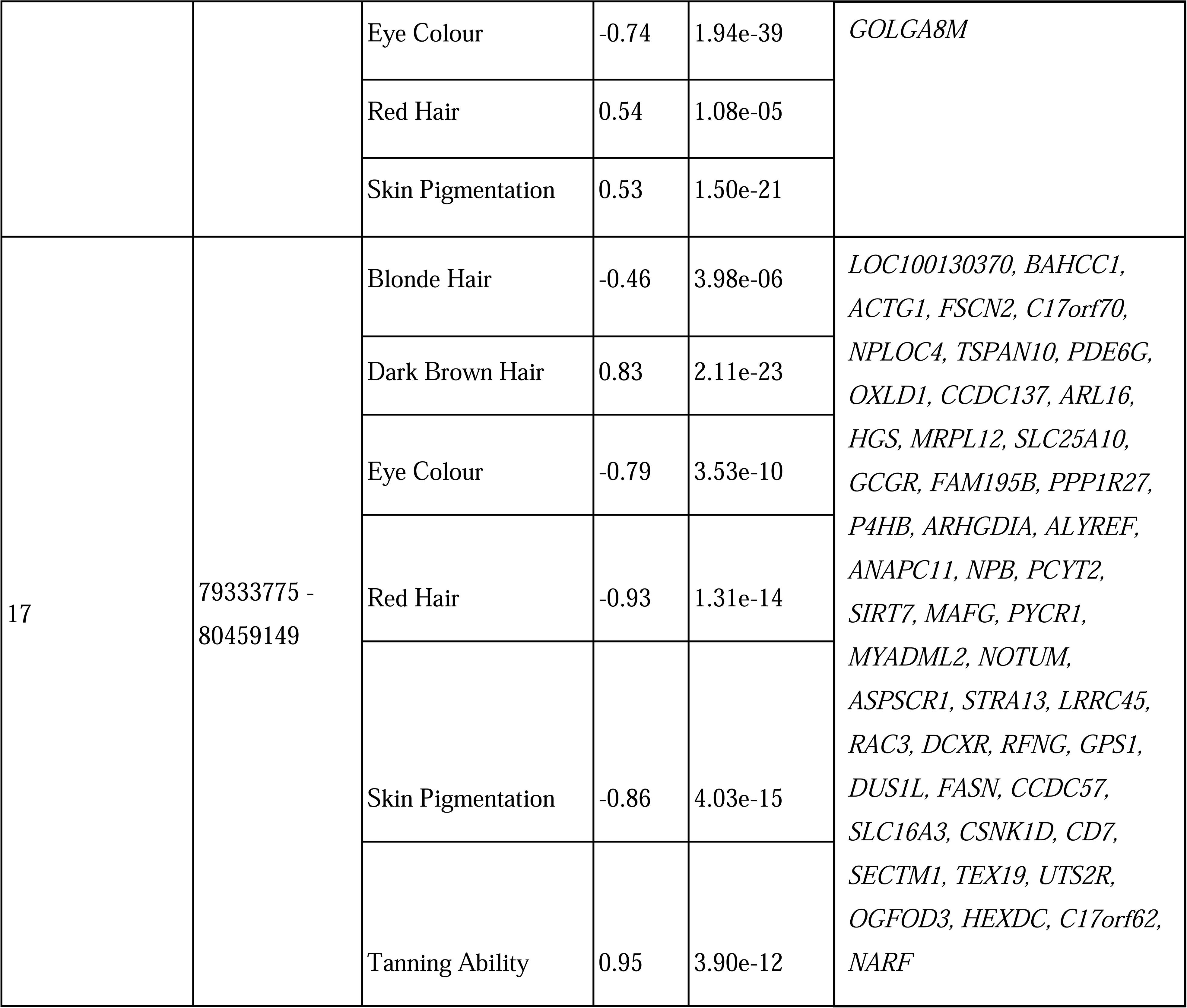
Statistically significant findings in the LAVA local correlation analyses between left eye overall average RPE thickness and other pigmentation-related traits. The genes listed are all these that, at least partially, overlap with the relevant locus. RPE, retinal pigment epithelium; RPS, retinal pigmentation score.

### Mendelian randomisation

Mendelian randomisation analysis could be conducted for 10,310 exposure traits. Three exposures from non-overlapping GWAS surpassed the multiple testing correction of the MRE IVW and were nominally significant in the weighted median, weighted mode and throughout leave one out analysis (Supplementary Table S4). These were serum apolipoprotein B (beta = -0.08, MRE IVW p = 7.07E-13), concentration of small LDL particles (beta = -0.07, MRE IVW p = 9.89E-15) and cathepsin H (beta = 0.05, MRE IVW p = 5.11E-32). The full results are presented in Supplementary Table S4.

## Discussion

We conducted phenotypic and genotypic analyses to advance our understanding of how pigmentation influences OCT-measured RPE thickness. This study is not concerned with the histological thickness of RPE and its relation to pigmentation, which may well differ given our results are most likely a product of the optical properties of the tissue. We found that RPE thickness varies according to fundus/choroidal pigmentation (RPS) and, to a lesser extent, according to skin colour. The latter variation was found to be driven by subjects with darker pigmentation (those with brown or black skin). Second, the GWAS that we conducted highlighted that genetic variation in pigmentation-related genes influences RPE thickness, with variants in *TYR*, *OCA2*-*HERC2*, *MREG*, and *TSPAN10* playing an important role. These include rs1126809 [*TYR*:c.1205G>A (p.Arg402Gln)], a missense variant in *TYR,* the gene that encodes tyrosinase, an enzyme which catalyses the conversion of tyrosine to melanin (Green *et al*., 2024). *TYR* is associated with a range of pigmentation-related phenotypes including albinism, hair colour, skin colour, ease of tanning and risk of skin cancer (Green et al., 2024; Lona-Durazo et al., 2019; Morgan et al., 2018; Rajesh et al., 2023; Simcoe et al., 2021). Another key signal was linked to rs12913832, a non-coding variant in a region near the *OCA2* and *HERC2* genes that has been identified as an enhancer regulating *OCA2*. *OCA2* encodes a transmembrane protein that functionally interacts with tyrosinase and the rs12913832 variant has been found to be a strong predictor of eye colour and associated with skin and hair colour (Lona-Durazo et al., 2019; Morgan et al., 2018; Simcoe et al., 2021). Additional lead variants associated with pigmentation included rs1800407 (OCA2) and rs3770526, which resides within *MREG,* the protein coding gene for melanoregulin, which is implicated in melanocyte differentiation and melanosome transport. Similar findings have been documented in GWAS of other pigmented phenotypes discussed in this paper including RPS, hair colour, skin colour, iris colour and tanning ability (Lona-Durazo et al., 2019; Morgan et al., 2018; Simcoe et al., 2021). Additionally, as shown in a recent GWAS variants in pigmentation genes including *TYR* and *HERC2/OCA2* are related to AMD risk. Interestingly, this study also found evidence suggesting that a range of pigmentation traits are causally related to AMD risk. These findings serve to highlight the critical roles of pigmentation in RPE physiology and fit into a broader literature supporting protective roles of ocular pigmentation against retinal degeneration (Gorman et al., 2024; Kauffman & Han, 2024). Notably, several of these pigmentation genes have previously been shown to influence the morphology of the vasculature and the thickness of non-pigmented layers, which appears likely to reflect the effect of pigmentation on the resolution of structures in optical scanning modalities. In our analysis, the optical properties of the tissue with respect to pigmentation are of direct relevance to the biology and therefore are central to the results rather than a confounding feature.

Notably, whilst there is highly significant phenotypic correlation between tanning ability, skin colour, hair colour and RPS, RPE thickness bears no significant phenotypic correlation to pigmentation-related traits other than RPS, and even then, there is only weak signal. We have also found that RPS, iris colour and RPE thickness are not genetically correlated with extraocular pigmentation-related traits on a global level. It is highlighted that RPE thickness and RPS were not found to share significant global genetic correlation with this finding being in keeping with the divergent pigmentation biology between the RPE and the choroid. This is likely underpinned by the differing embryological origins and physiology of RPE and uveal melanocytes. When exploring the local genetic relationships between RPE and other pigmented tissues, we found that there were several regions of local genetic correlation in loci containing genes related to pigmentation including *TYR, RAB38, OCA2-HERC2* and *TSPAN10*. We also found that there were substantially more significant local genetic correlations between extraocular pigmented tissues than ocular ones (Figure 6). These results suggest that the genetic determinants of pigmentation of ocular and extraocular tissues differ with the exception of a small number of regions containing influential genes that have a role in melanosome function. This differs to extraocular traits which are significantly correlated with each other on a global genetic level, and in a more diverse range of local genetic regions (as can be seen in Supplementary Table 3).

It is noted that this study also highlighted non-pigment-related determinants of RPE thickness. First, the conducted GWAS identified variants known to be associated with ophthalmic diseases and OCT features, including rs1410996 and rs443134 (*CFHR3*) which are associated with age-related macular degeneration; rs57819090 (*C8orf74*) and rs62075723 (*NPLOC4-TSPAN10*) which are associated with retinal nerve fibre layer thickness; and rs2625955 and rs4840499 which are within the protein coding gene *RHO*, which encodes the primary photoreceptor molecule of vision (rhodopsin) which resides within rod cells of the retina (Lenahan *et al*., 2020). Further, the phenome-wide Mendelian randomisation analysis that we conducted identified causal roles for two lipid traits (serum apolipoprotein B and concentration of small LDL particles) and cathepsin H as determinants of RPE thickness. Numerous lipid traits were nominally significant in relation to RPE thickness but did not pass a multiple testing correction (Supplementary Table 4). Because lipid traits have a similar genetic architecture, there is a multicollinearity issue which precludes a statement that any particular lipid-related trait is uniquely causally related to RPE thickness (Julian, Cooper-Knock, *et al*., 2023). Instead, this result simply highlights the critical role that lipid homeostasis plays in RPE physiology, as has been documented in previous studies of RPE biology as well as in studies of age-related macular degeneration risk (Julian, Cooper-Knock, *et al*., 2023). Cathepsins are lysosomal proteases, with functions in the RPE documented for Cathepsin D and S which are involved in the degradation of photoreceptor outer segments (Im & Kazlauskas, 2007; Julian, Cooper-Knock, et al.,

2023). Previously, in a phenome-wide causal inference study of AMD, we found that Cathepsin F was causally related to AMD risk (Gorman et al., 2024; Julian, Cooper-Knock, et al., 2023). Here, we found that serum Cathepsin H is causally associated with greater RPE thickness. Although, to our knowledge, cathepsin H has not previously been described in direct relation to RPE physiology, the coding gene *CTSH* is expressed in the retina and so could plausibly have important roles (Senabouth *et al*., 2022). Cathepsin H has recognised roles in the complement cascade, in which it is one of the few non-complement proteases which cleaves C5 to potent chemotaxin C5a (Wang *et al*., 2023). Plausibly, the role of Cathepsin H in complement could be relevant to its influence over RPE thickness given the recognised role of complement over RPE physiology and pathophysiology. Finally, whilst we are not aware of studies linking Cathepsin H to pigmentation, there are clearly documented roles for other cathepsins in melanosome degradation. For instance, Cathepsin V has been directly implicated in melansome degradation and is differentially expressed according to skin pigmentation while Cathepsin L has been shown to influence proliferation, differentiation and apoptosis of hair follicle melanocytes (Homma et al., 2018; Kim et al., 2021; Tobin et al., 2002). Therefore, there may also be merit in future studies exploring the roles of cathepsins in pigmentation of the RPE.

In conclusion, this study provides insights into the role of pigmentation in OCT-measured RPE thickness, highlights regions of local genetic correlation between RPE thickness and other pigmentation-related traits in blocks containing pigmentation genes, and identifies causal roles for lipid homeostasis and Cathepsin H levels in RPE thickness.

## Acknowledgements

We acknowledge the contribution of the UK Biobank Eye and Vision Consortium. Members of this consortium include: Naomi Allen, Tariq Aslam, Denize Atan, Sarah Barman, Jenny Barrett, Paul Bishop, Graeme Black, Tasanee Braithwaite, Roxana Carare, Usha Chakravarthy, Michelle Chan, Sharon Chua, Alexander Day, Parul Desai, Bal Dhillon, Andrew Dick, Alexander Doney, Cathy Egan, Sarah Ennis, Paul Foster, Marcus Fruttiger, John Gallacher, David Garway-Heath, Jane Gibson, Jeremy Guggenheim, Chris Hammond, Alison Hardcastle, Simon Harding, Ruth Hogg, Pirro Hysi, Pearse Keane, Peng Tee Khaw, Anthony Khawaja, Gerassimos Lascaratos, Thomas Littlejohns, Andrew Lotery, Robert Luben, Phil Luthert, Tom Macgillivray, Sarah Mackie, Savita Madhusudhan, Bernadette Mcguinness, Gareth Mckay, Martin Mckibbin, Tony Moore, James Morgan, Eoin O’Sullivan, Richard Oram, Chris Owen, Praveen Patel, Euan Paterson, Tunde Peto, Axel Petzold, Nikolas Pontikos, Jugnoo Rahi, Alicja Rudnicka, Naveed Sattar, Jay Self, Panagiotis Sergouniotis, Sobha Sivaprasad, David Steel, Irene Stratton, Nicholas Strouthidis, Cathie Sudlow, Zihan Sun, Robyn Tapp, Dhanes Thomas, Emanuele Trucco, Adnan Tufail, Ananth Viswanathan, Veronique Vitart, Mike Weedon, Alastair K. Denniston, Peng T. Khaw, Konstantinos Balaskas, Cathy Williams, Katie Williams, Jayne Woodside, Max Yates, Jennifer Yip, Yalin Zheng.

## Author Contribution Statement

T.H.J completed the analysis presented in this article, with methodological input from the other authors. All authors were involved in study design and manuscript drafting.

## Declaration of Interests

E.B. is a paid consultant and equity holder of Oxford Nanopore, a paid consultant to Dovetail and a non-executive director of Genomics England, a limited company wholly owned by the UK Department of Health and Social Care. All other authors declare no competing interests.

## Ethical Approval

This study was conducted using UK Biobank data. UK Biobank has approval from the North West Multi-centre Research Ethics Committee (MREC) as a Research Tissue Bank (RTB) approval.

## Data availability

The summary statistics from our genome wide association study will be made publicly available upon publication. All other generated statistics have been presented within the supplementary content of this article.

## Funding Statement

We acknowledge the following sources of funding: the Medical Research Council (MRC) (MR/Z504105/1, Clinical Research Training Fellowship to T.H.J.); the Wellcome Trust (224643/Z/21/Z, Clinical Research Career Development Fellowship to P.I.S.); the UK National Institute for Health Research (NIHR) Clinical Lecturer Programme (CL2017-06-001 to P.I.S.); the NIHR Manchester Biomedical Research Centre (NIHR 203308 to P.I.S.); the NIHR Academic Clinical Fellowship (NIHR ACF-2021-06-011 to T.H.J); and the EMBL European Bioinformatics Institute (EMBL-EBI) (E.B., T.F.). The UK Biobank Eye and Vision Consortium is supported by funding from the NIHR Biomedical Research Centre at Moorfields Eye Hospital and UCL Institute of Ophthalmology, the Alcon Foundation and the Desmond Foundation.

## Supporting Information Captions

**Supplementary Table 1**: the results of phenotypic correlation analysis using Kendall’s Tau. The results are presented as Kendall’s Tau coefficient followed by an indicator of p value. * = p<0.05, ** = 0<0.01, *** = p<0.001

**Supplementary Table 2**: The results of pairwise comparisons using Wilcoxon rank sum test with continuity correction, testing for significant differences in RPE thickness and RPS according to skin colour.

**Supplementary table 3a**: A table detailing only multiple testing corrected significant bivariate relationships between pigmented traits. Locus number is an arbitrary figure assigned at the start of the analysis. Start and stop are the base pare locations at the start and end of the defined locus. N SNPs is the number of SNPs in that locus. Rho is the measure of local regression. Upper and lower values represent the confidence intervals. r2 is the proportion of genetic signal explained jointly by the predictors. The genes detailed are those contained within the locus, and the figure in brackets in this column is the proportion of overlap between the gene and the locus. N PCs is the number of principal components.

**Supplementary table 3b**: A table detailing all tested bivariate relationships between pigmented traits. Locus number is an arbitrary figure assigned at the start of the analysis. Start and stop are the base pare locations at the start and end of the defined locus. N SNPs is the number of SNPs in that locus. Rho is the measure of local regression. Upper and lower values represent the confidence intervals. r2 is the proportion of genetic signal explained jointly by the predictors. N PCs is the number of principal components.

**Supplementary table 3c:** A table detailing all tested univariate local genetic heritability of traits at each locus. Locus number is an arbitrary figure assigned at the start of the analysis. Start and stop are the base pare locations at the start and end of the defined locus. N SNPs is the number of SNPs in that locus. H2 is the heritability estimate. N PCs is the number of principal components.

**Supplementary table 4:** A table detailing all MR analyses conducted. Trait ID is the identifier of the trait in the Two Sample MR R package. Trait name is the full name of the trait. nSNP is the number of exposure SNPs in the analysis. Inclusion p value is the p value threshold at which SNPs for the exposure trait were selected. OR is odds ratio. LCI95 and UCI95 are the lower and upper 95% confidence intervals respectively. MRE IVW is the multiplicative random effects inverse variance weighted analysis. SE is standard error. LOO is leave one out analysis, which states if the analysis was significant throughout an MRE IVW leave one out analysis. The pleiotropic SNP removal and outlier SNP removal columns indicate whether any SNPs had to be removed during Steiger filtering or by radial MR outlier detection. Reverse MR analyses with the IVW are presented only where the primary analysis was significant. Only analyses detailed in the manuscript have been manually inspected for sample overlap, and it is essential that readers check for sample overlap before drawing conclusions regarding causation from this data, as substantial sample overlap biases the analysis toward a significant causal relationship.

**Supplementary Figure 1:** A histogram which illustrates the spread of the left eye overall RPE thickness in microns.

**Supplementary Figure 2:** A histogram which illustrates the spread of the right eye overall RPE thickness in microns.

**Supplementary Figure 3:** A histogram which illustrates the spread of the left eye overall RPE thickness in the replication cohort, in microns.

**Supplementary Figure 4:** A histogram which illustrates the spread of the right eye central RPE thickness in microns.

**Supplementary Figure 5:** A histogram which illustrates the spread of the left eye centre RPE thickness in microns.

**Supplementary Figure 6:** A histogram which illustrates the spread of the right eye mean inner RPE thickness in microns.

**Supplementary Figure 7:** A histogram which illustrates the spread of the left eye mean inner RPE thickness in microns.

**Supplementary Figure 8:** A histogram which illustrates the spread of the left eye mean outer RPE thickness in microns.

**Supplementary Figure 9:** A histogram which illustrates the spread of the right eye mean outer RPE thickness in microns.

**Supplementary Figure 10:** A histogram which illustrates the spread of the left eye retinal pigmentation score.

**Supplementary Figure 11:** A scatter plot indicating that there is very little correlation between RPE thickness and RPS. The red line is a line of best fit.

**Supplementary Figure 12:** A density plot which shows the distribution of RPE thickness according to self-reported skin colour.

**Supplementary Figure 13:** A density plot which shows the distribution of retinal pigmentation score according to self-reported skin colour.

**Supplementary Figure 14:** A density plot which shows the distribution of RPE thickness according to self-reported ethnicity.

**Supplementary Figure 15:** A density plot which shows the distribution of retinal pigmentation score according to self-reported ethnicity.

**Supplementary Figure 16:** A beta-beta plot showing the correlation between single nucleotide polymorphisms (SNPs) betas which were genome wide significant in our discovery GWAS, and their beta values in the replication GWAS.

**Supplementary Figure 17:** The Manhattan plot for our GWAS of left eye overall RPE thickness. Genome-wide significant SNPs are annotated.

**Supplementary Figure 18:** The QQ plot for our GWAS of left eye overall RPE thickness.

**Supplementary Figure 19:** The Manhattan plot for our GWAS of right eye overall RPE thickness. Genome-wide significant SNPs are annotated.

**Supplementary Figure 20:** The QQ plot for our GWAS of right eye overall RPE thickness.

**Supplementary Figure 21:** The Manhattan plot for our GWAS of left eye centre RPE thickness. Genome-wide significant SNPs are annotated.

**Supplementary Figure 22:** The QQ plot for our GWAS of left eye centre RPE thickness.

**Supplementary Figure 23:** The Manhattan plot for our GWAS of right eye centre RPE thickness. Genome-wide significant SNPs are annotated.

**Supplementary Figure 24:** The QQ plot for our GWAS of right eye centre RPE thickness.

**Supplementary Figure 25:** The Manhattan plot for our GWAS of left eye mean inner RPE thickness. Genome-wide significant SNPs are annotated.

**Supplementary Figure 26:** The QQ plot for our GWAS of left eye mean inner RPE thickness.

**Supplementary Figure 27:** The Manhattan plot for our GWAS of right eye mean inner RPE thickness. Genome-wide significant SNPs are annotated.

**Supplementary Figure 28:** The QQ plot for our GWAS of right eye mean inner RPE thickness.

**Supplementary Figure 29:** The Manhattan plot for our GWAS of left eye mean outer RPE thickness. Genome-wide significant SNPs are annotated.

**Supplementary Figure 30:** The QQ plot for our GWAS of left eye mean outer RPE thickness.

**Supplementary Figure 31:** The Manhattan plot for our GWAS of right eye mean outer RPE thickness. Genome-wide significant SNPs are annotated.

**Supplementary Figure 32:** The QQ plot for our GWAS of right eye mean outer RPE thickness.

**Supplementary Figure 33:** The Manhattan plot for our GWAS of replication study of left eye overall RPE thickness.

**Supplementary Figure 34:** The QQ plot for our GWAS of replication study of left eye overall RPE thickness.

**Supplementary Figure 35:** Network chart detailing the local genetic correlations at 11:88359288-89209738, the locus containing *GABRG3, OCA2-HERC2, LOC645202, HERC2P11* and *GOLGA8M*. Blue connections indicate positive correlations whilst red indicates negative correlations. The opacity of the connection is greatest for weaker correlations in terms of coefficient, with stronger correlations indicated by wider, solid lines. All connections represent multiple testing corrected significant local genetic correlations.

**Supplementary Figure 36:** Network chart detailing the local genetic correlations at 11:88359288-89209738, the locus containing *GRM5, TYR* and *NOX4*. Blue connections indicate positive correlations whilst red indicates negative correlations. The opacity of the connection is greatest for weaker correlations in terms of coefficient, with stronger correlations indicated by wider, solid lines. All connections represent multiple testing corrected significant local genetic correlations.

## Notes

### Competing Interest Statement

The authors have declared no competing interest.

### Summary of Updates

The replication study omitted some eligible subjects and has been updated upon identification of this.

