## Supplementary Figures for "Pigmentation and retinal pigment epithelium thickness: a study of the phenotypic and genotypic relationships between ocular and extraocular pigmented tissues"

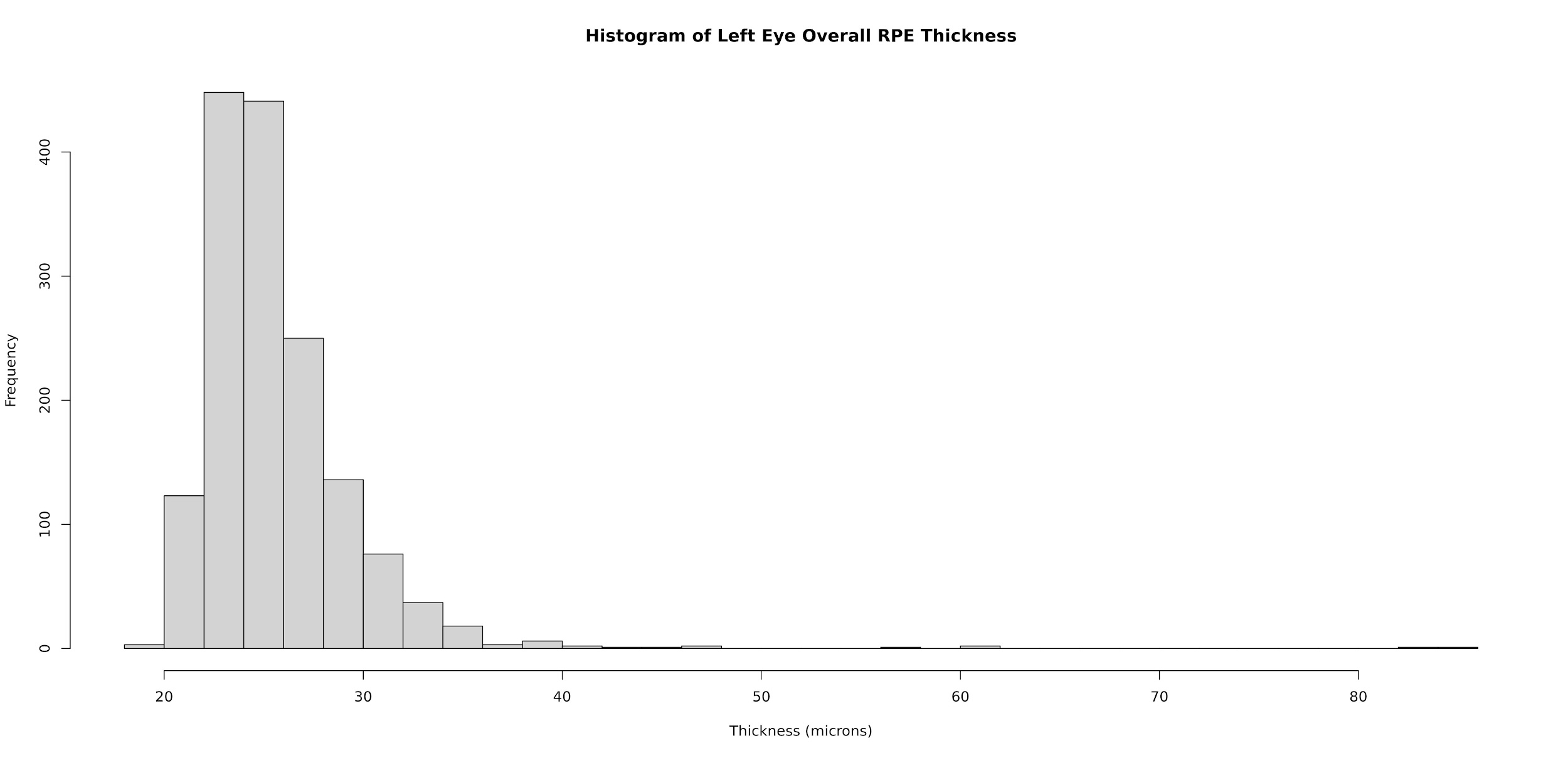
**Supplementary Figure 1:** A histogram which illustrates the spread of the left eye overall RPE thickness in microns.
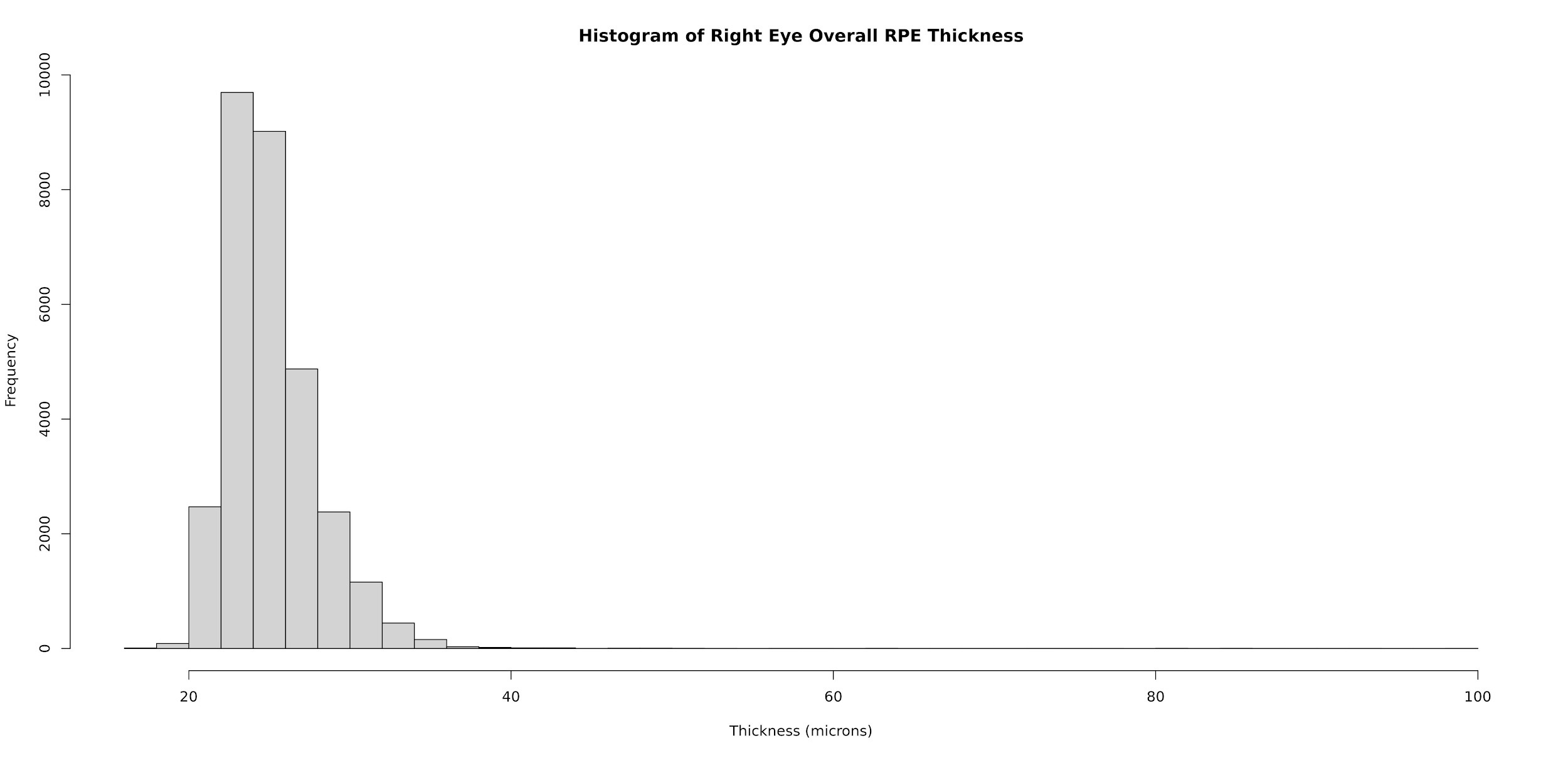
 **Supplementary Figure 2:** A histogram which illustrates the spread of the right eye overall RPE thickness in microns.
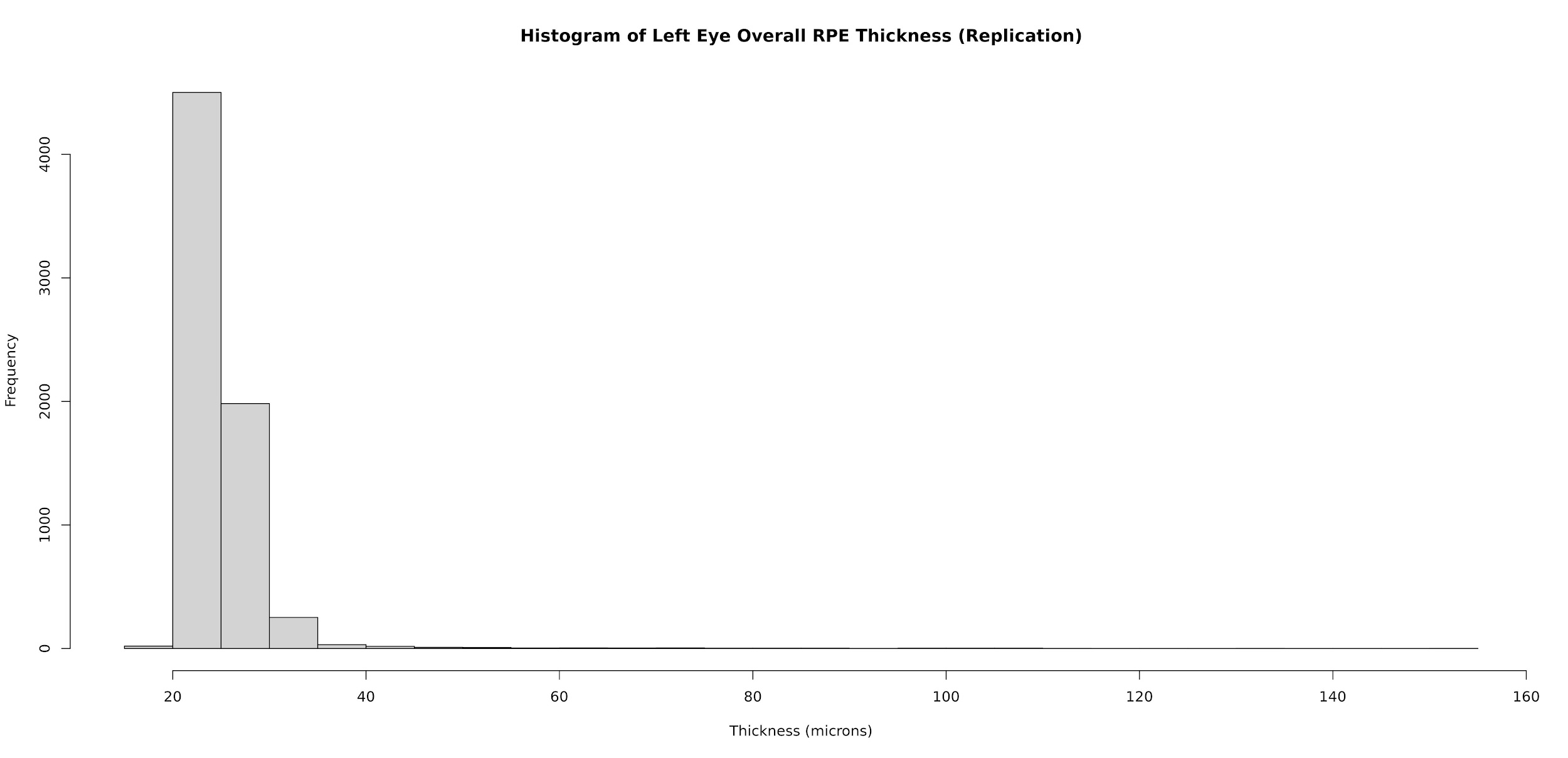
**Supplementary Figure 3:** A histogram which illustrates the spread of the left eye overall RPE thickness in the replication cohort, in microns.
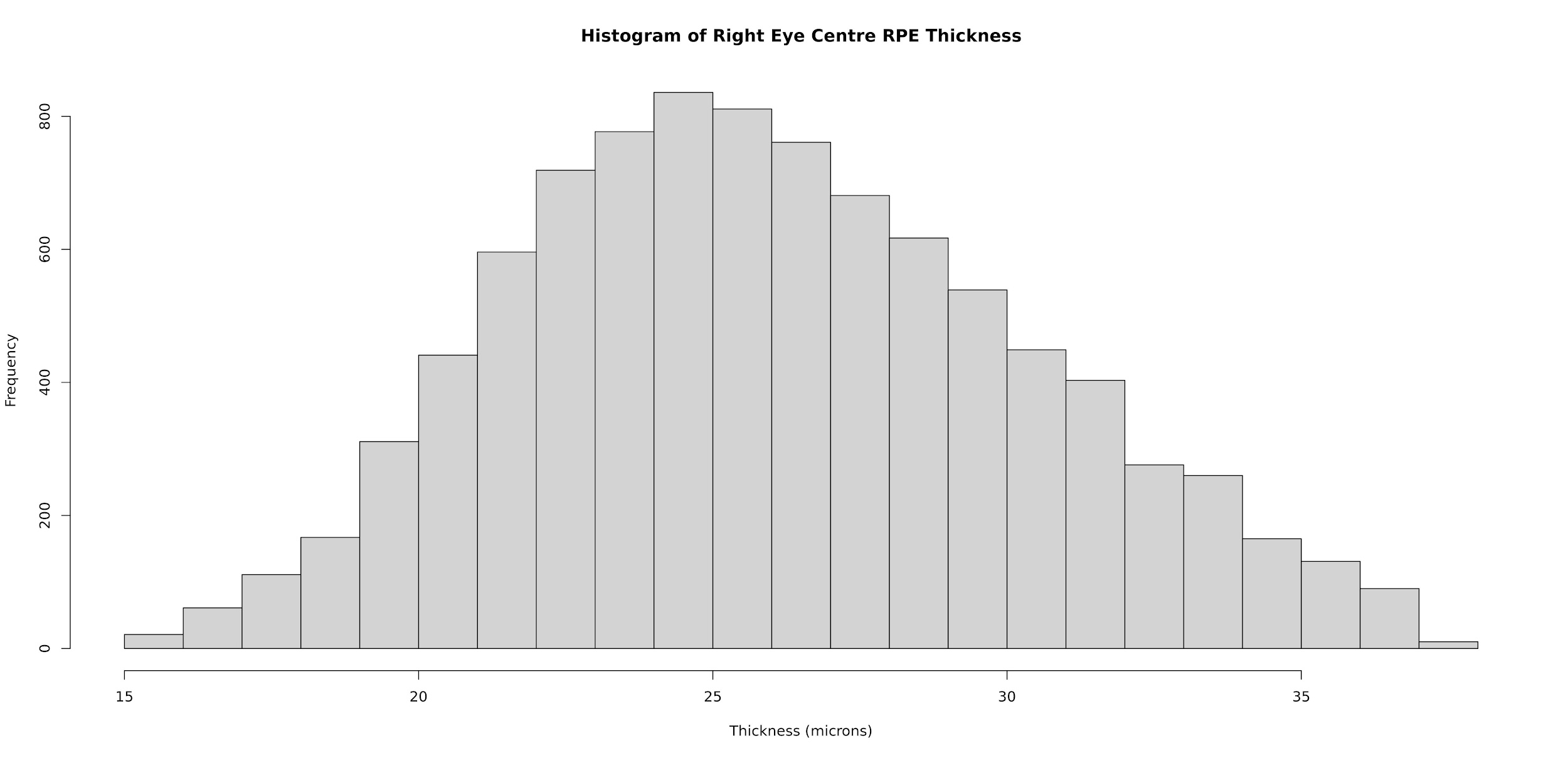
**Supplementary Figure 4:** A histogram which illustrates the spread of the right eye central RPE thickness in microns.
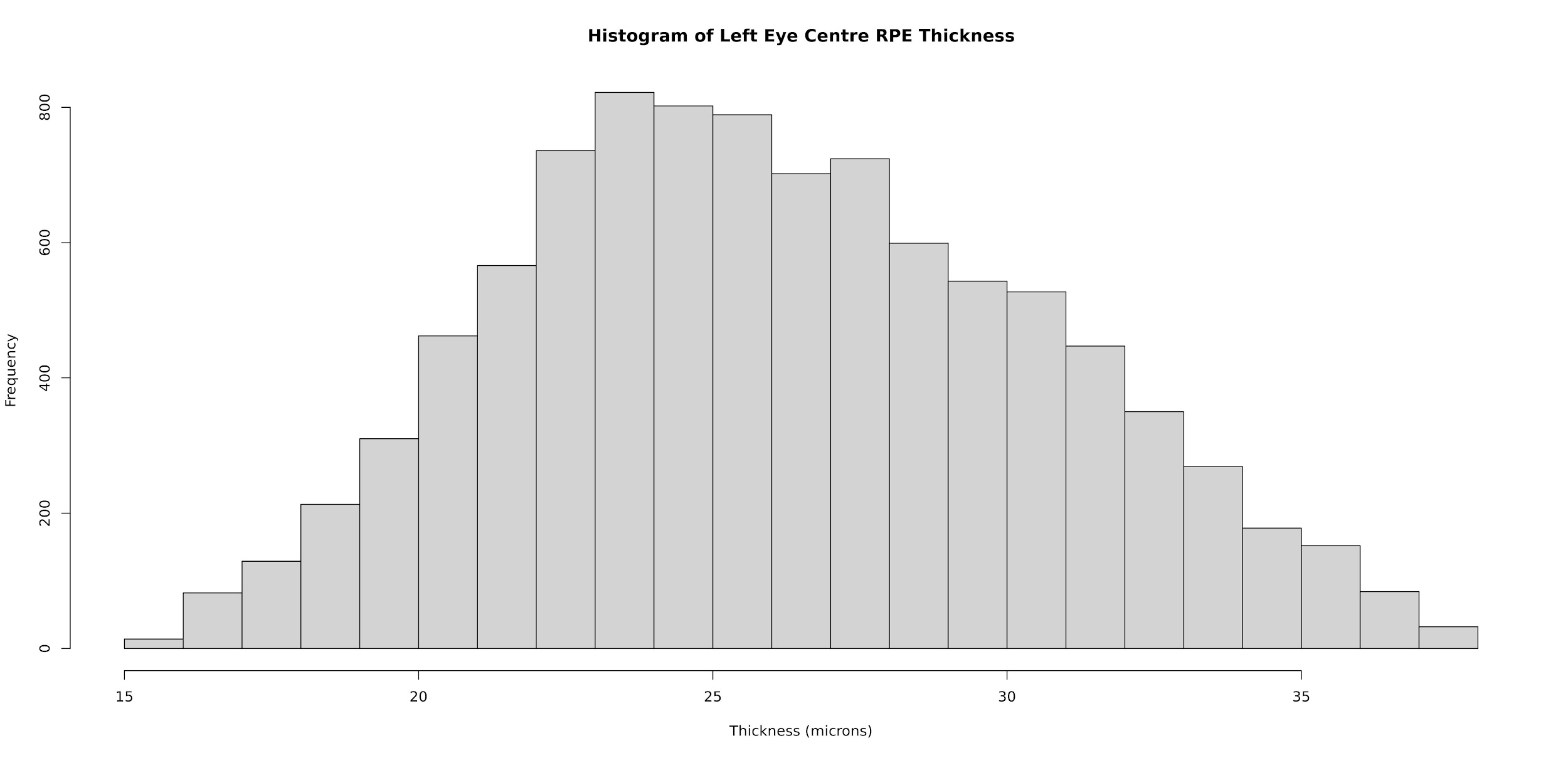
**Supplementary Figure 5:** A histogram which illustrates the spread of the left eye centre RPE thickness in microns.
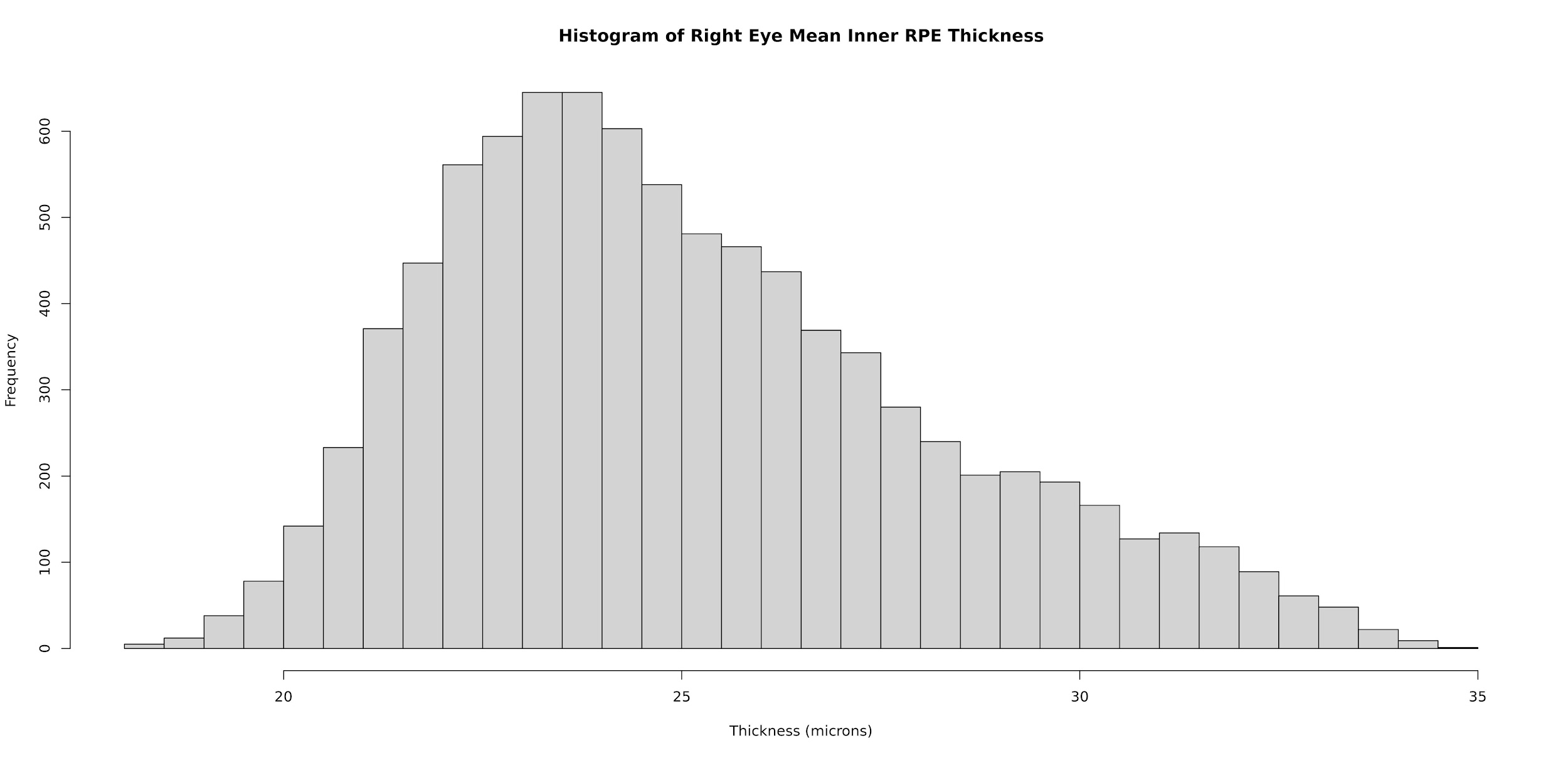
**Supplementary Figure 6:** A histogram which illustrates the spread of the right eye mean inner RPE thickness in microns.
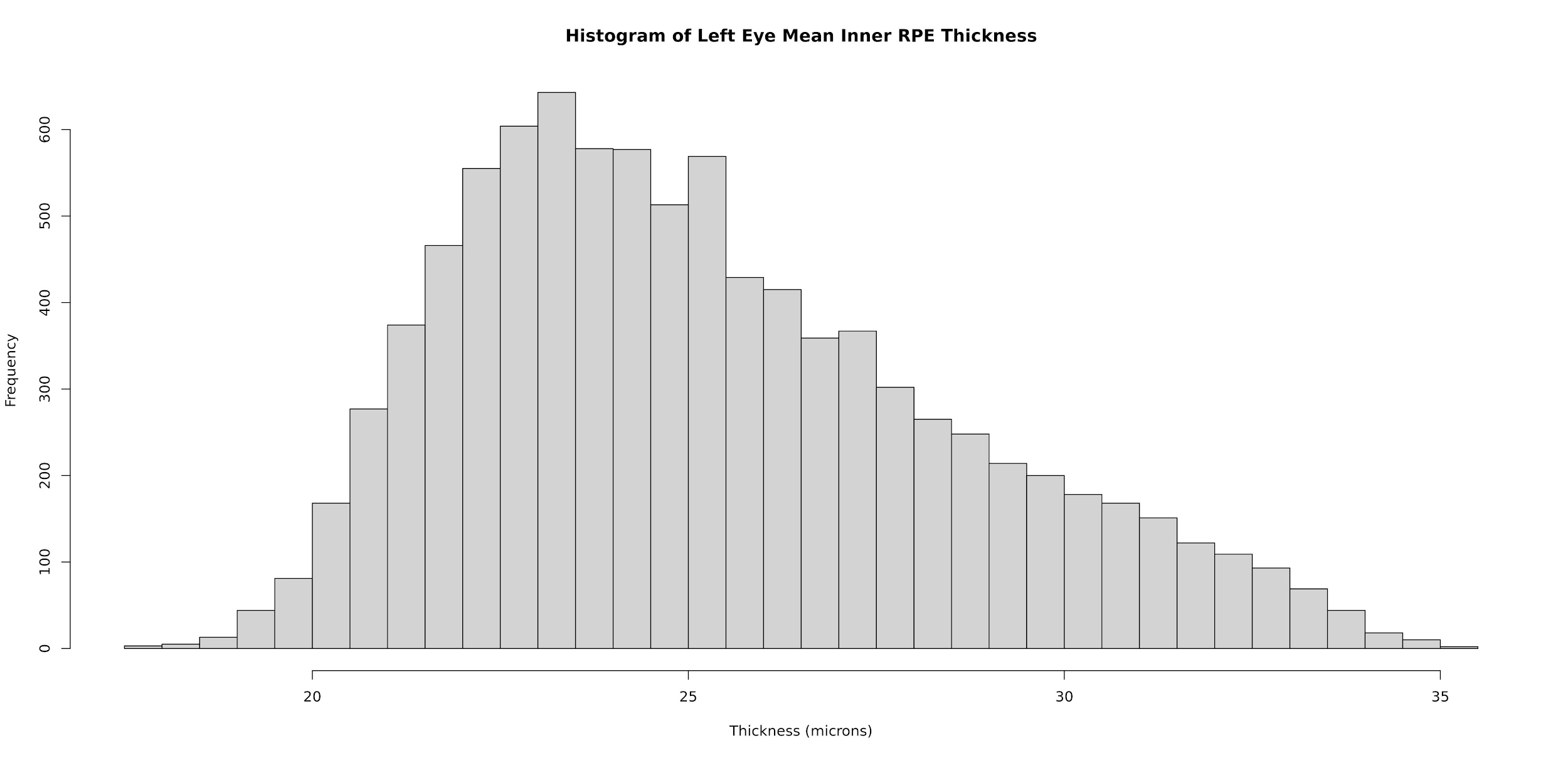
**Supplementary Figure 7:** A histogram which illustrates the spread of the left eye mean inner RPE thickness in microns.
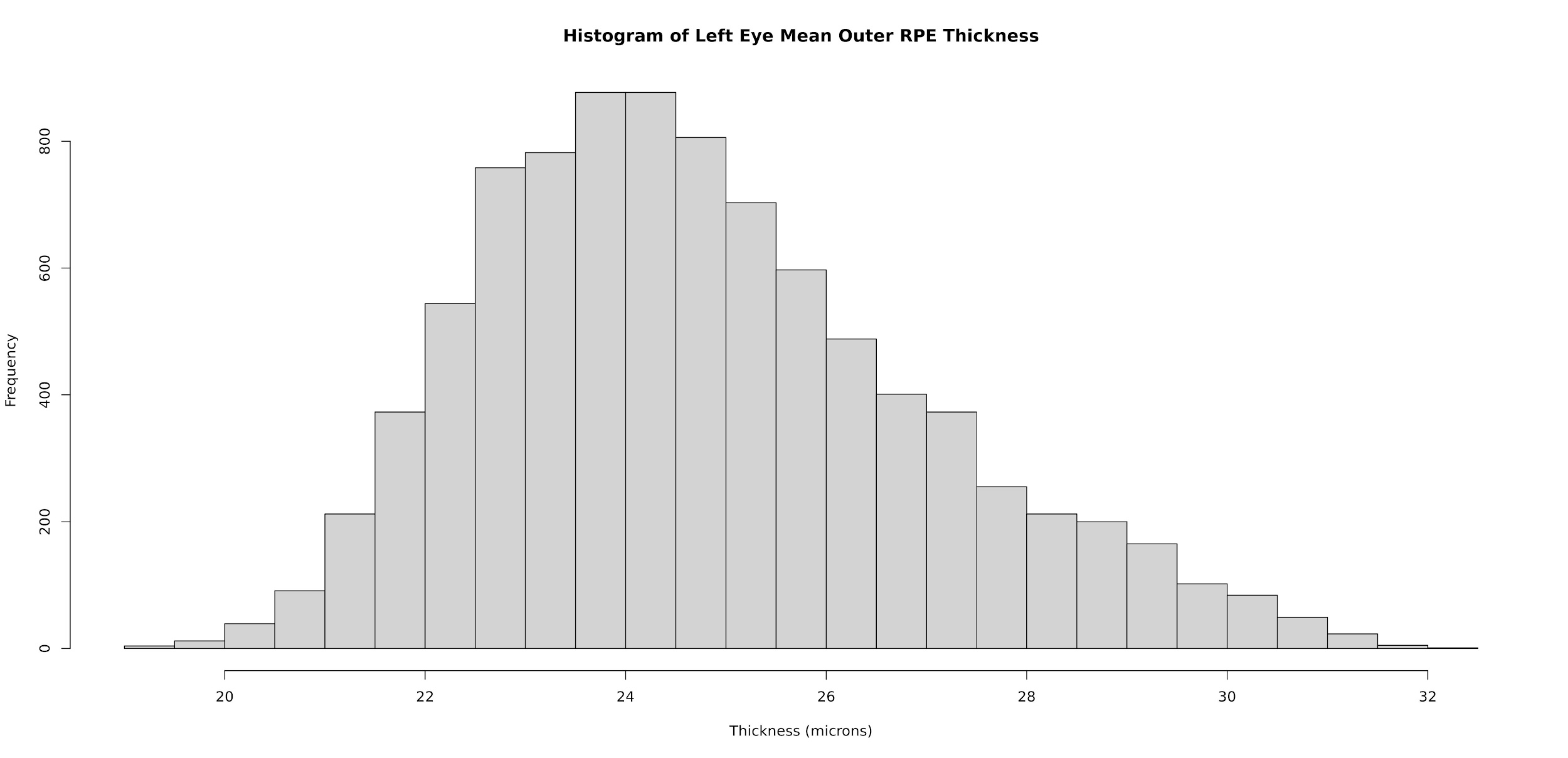
**Supplementary Figure 8:** A histogram which illustrates the spread of the left eye mean outer RPE thickness in microns.
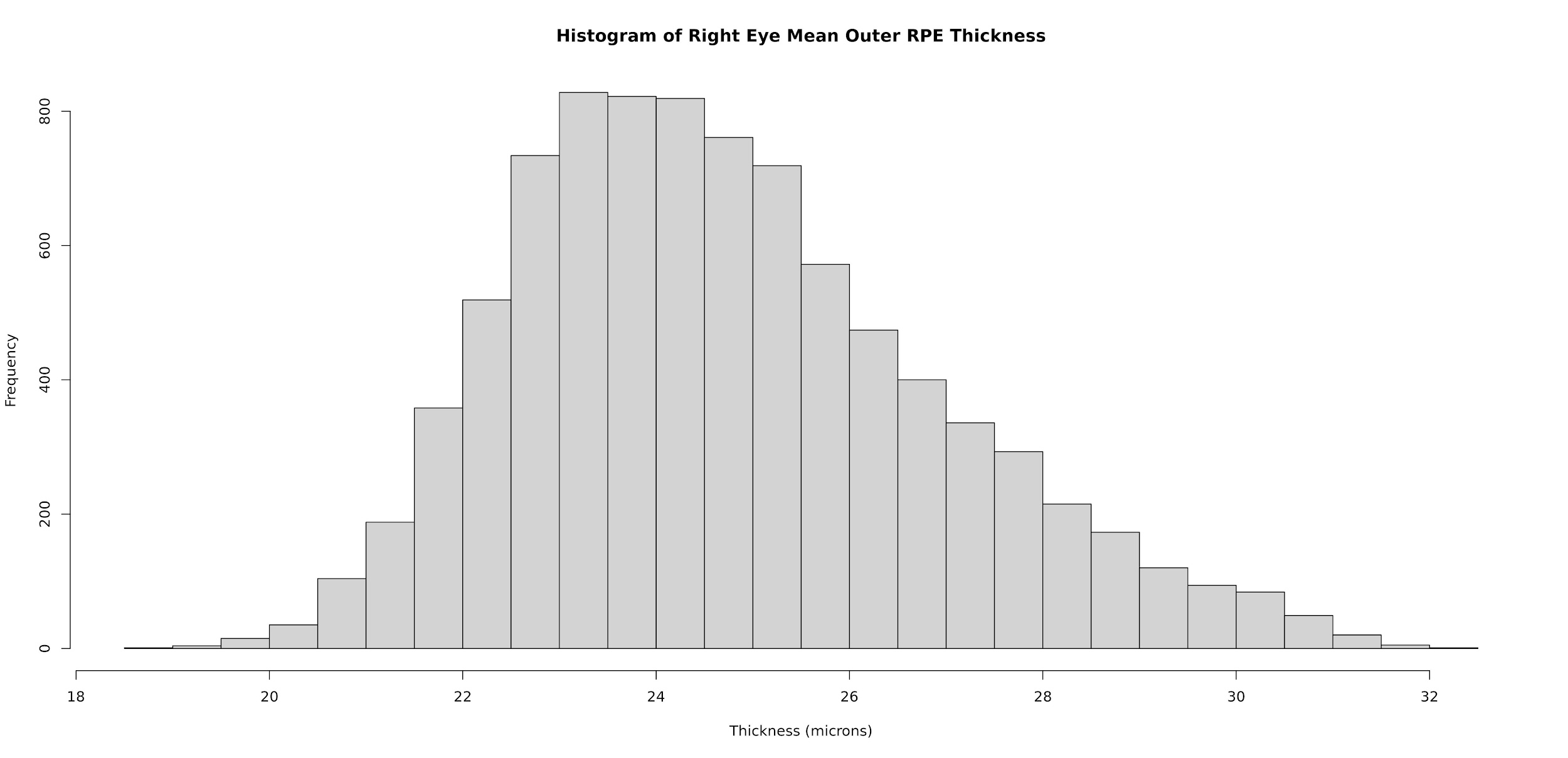
**Supplementary Figure 9:** A histogram which illustrates the spread of the right eye mean outer RPE thickness in microns.
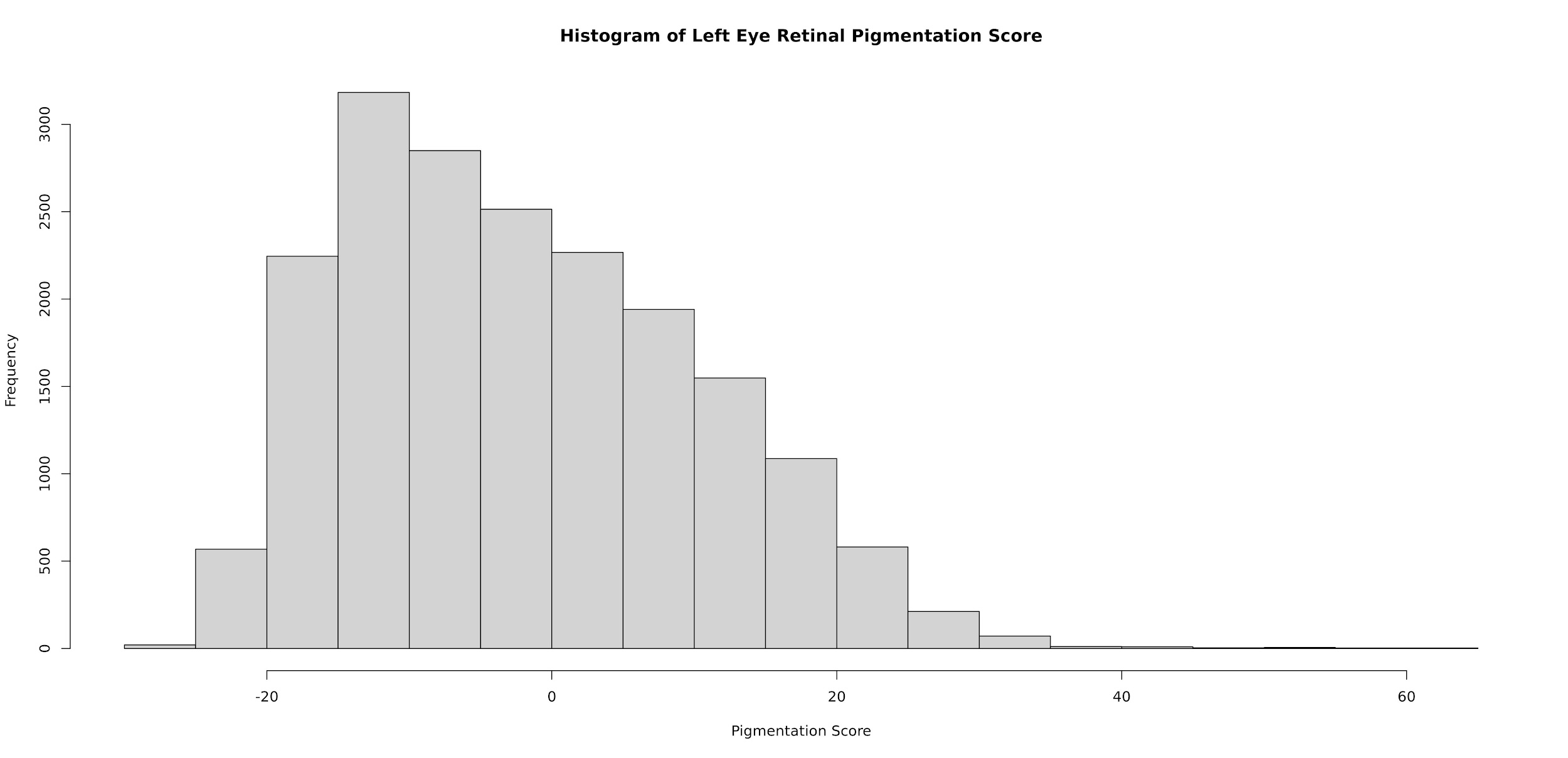
**Supplementary Figure 10:** A histogram which illustrates the spread of the left eye retinal pigmentation score.

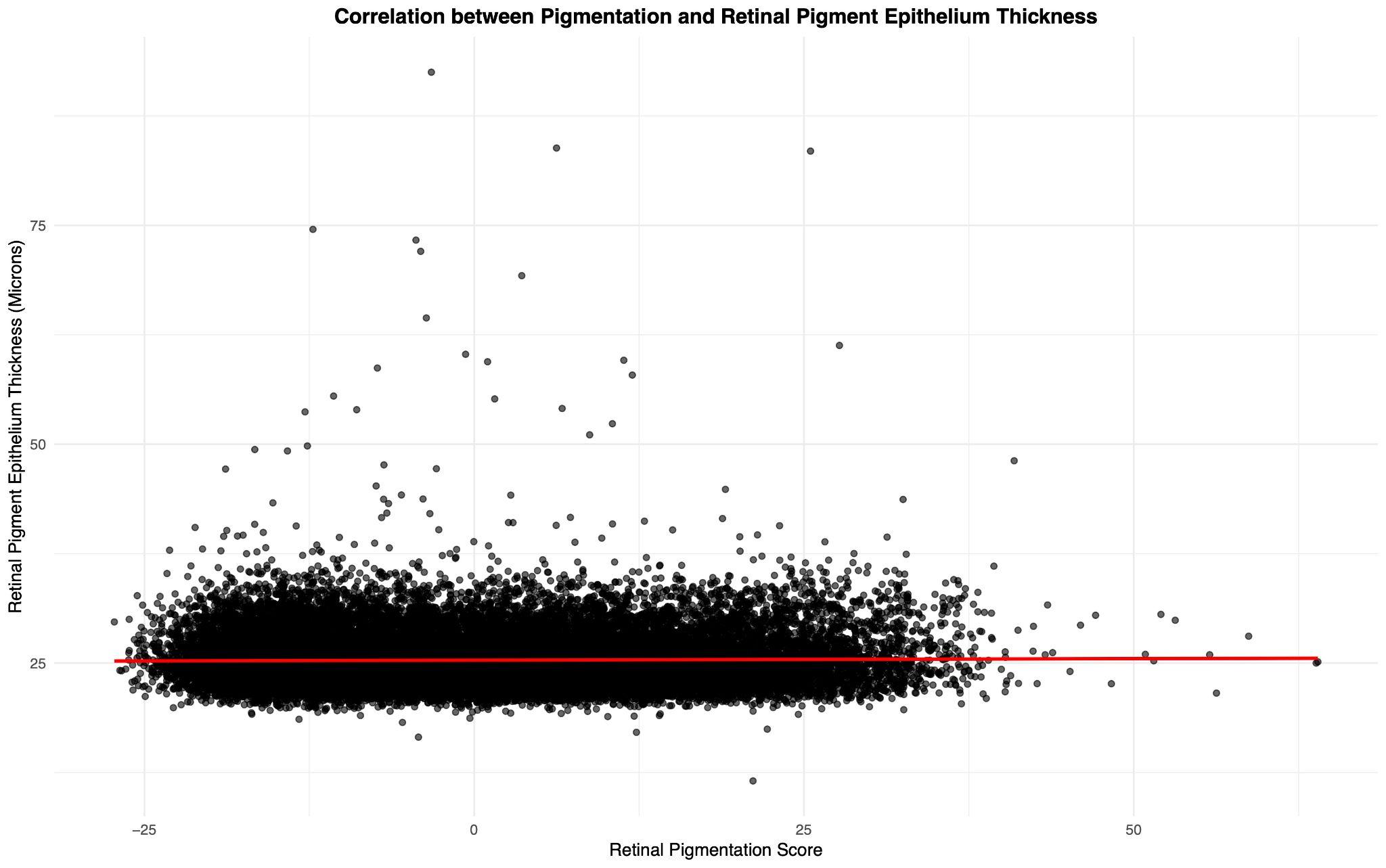
**Supplementary Figure 11:** A scatter plot indicating that there is very little correlation between RPE thickness and RPS. The red line is a line of best fit.

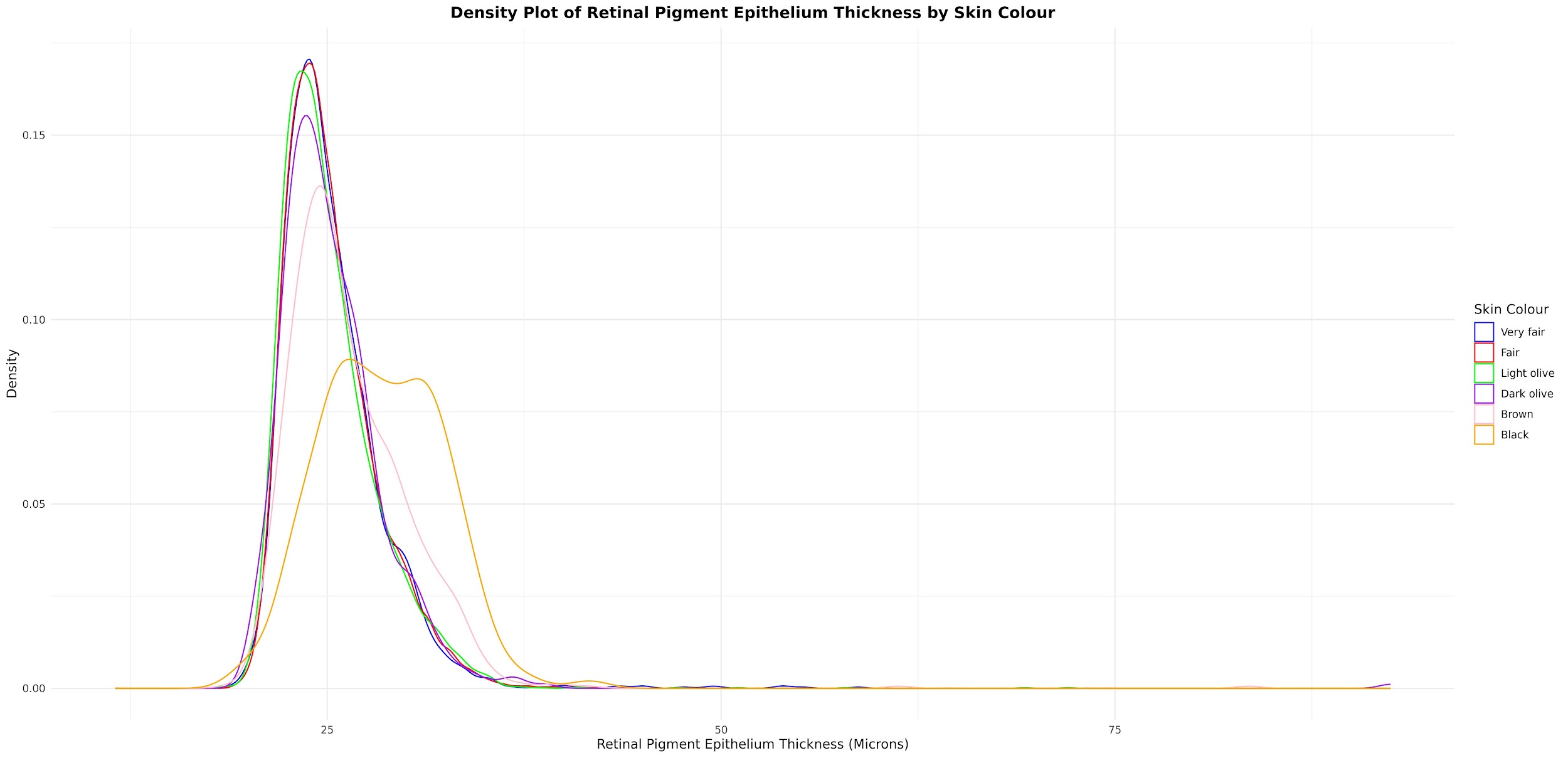
**Supplementary Figure 12:** A density plot which shows the distribution of RPE thickness according to self-reported skin colour.
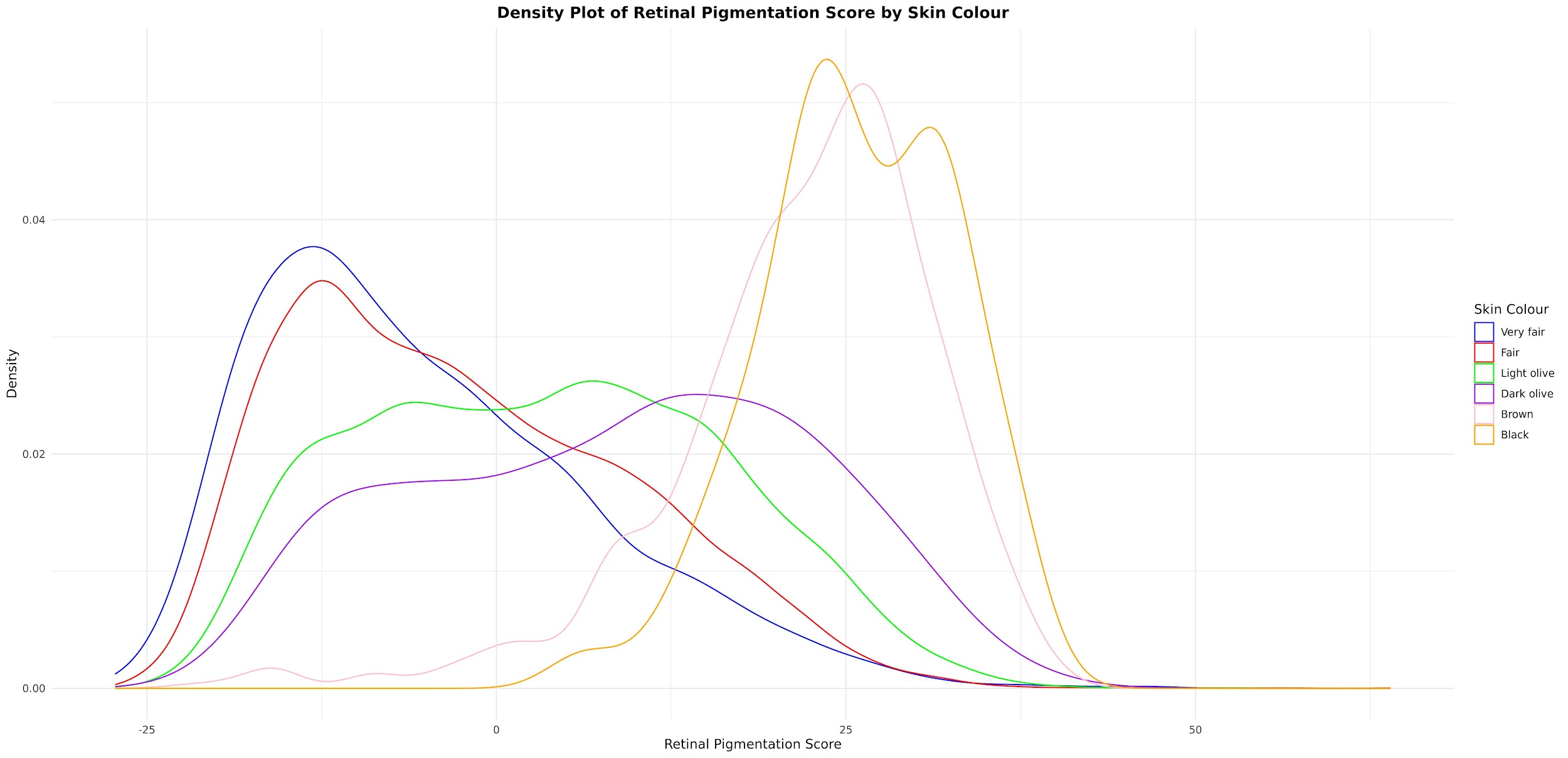
**Supplementary Figure 13:** A density plot which shows the distribution of retinal pigmentation score according to self-reported skin colour.
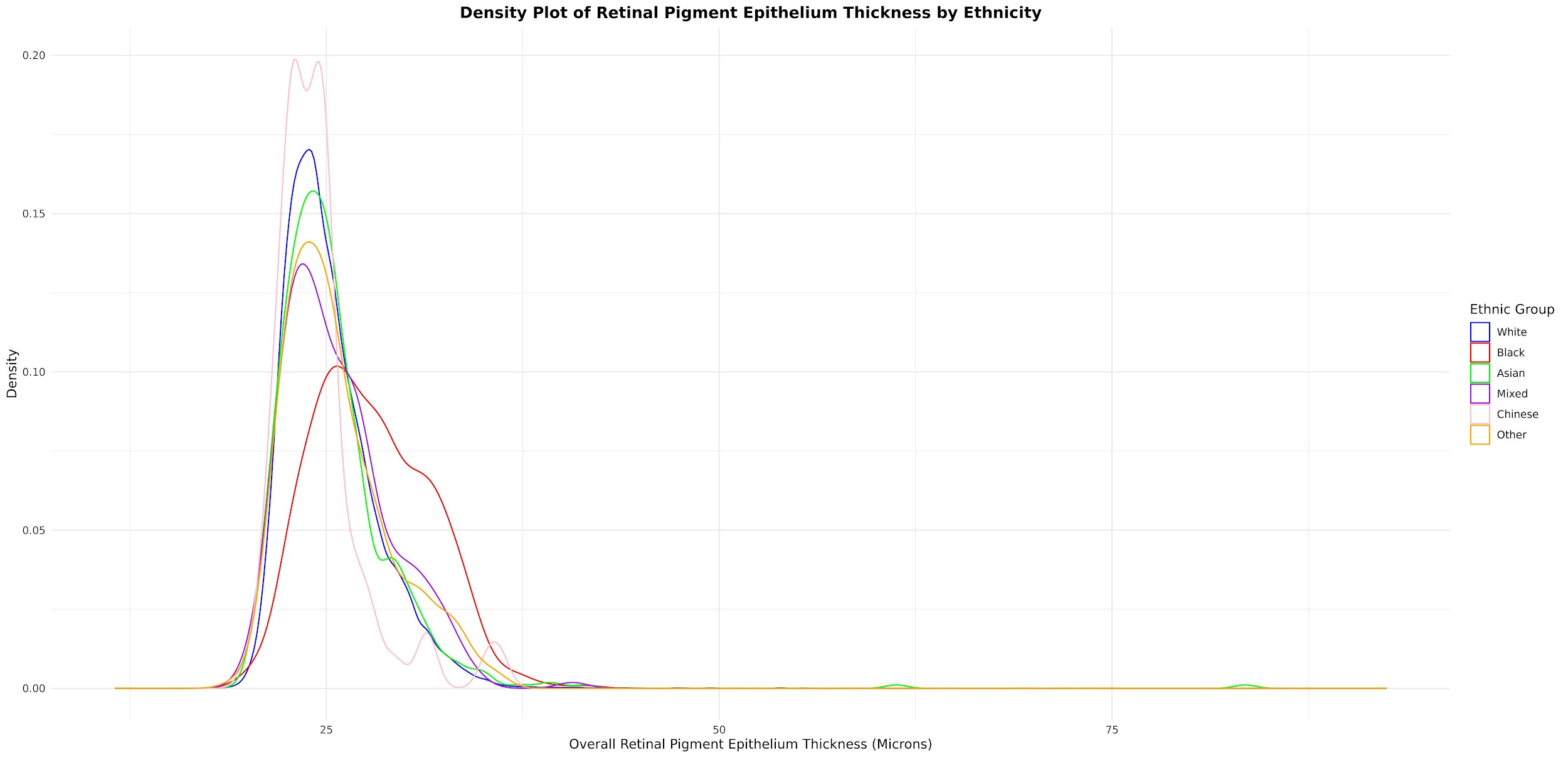
**Supplementary Figure 14:** A density plot which shows the distribution of RPE thickness according to self-reported ethnicity.
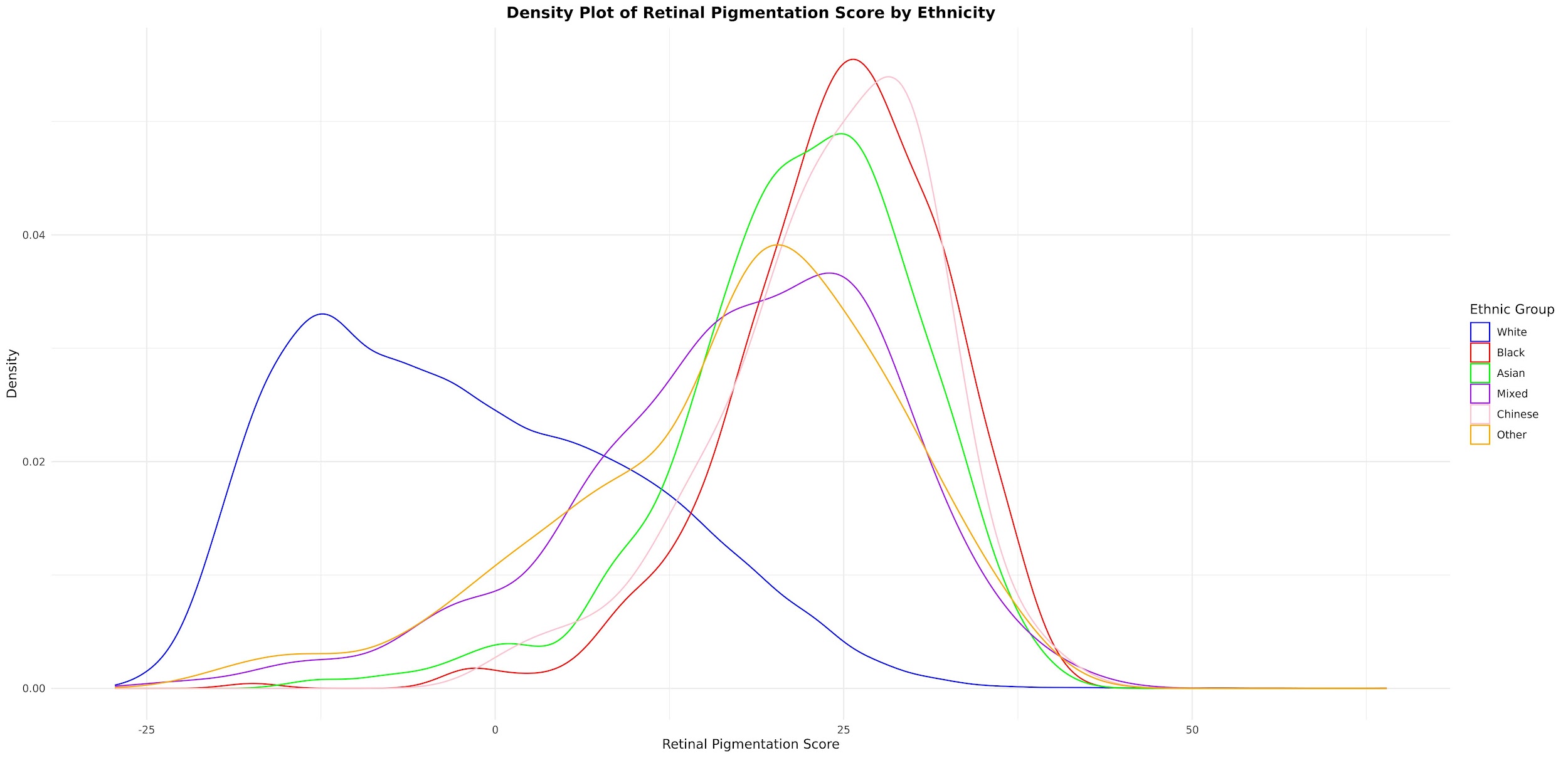
**Supplementary Figure 15:** A density plot which shows the distribution of retinal pigmentation score according to self-reported ethnicity.

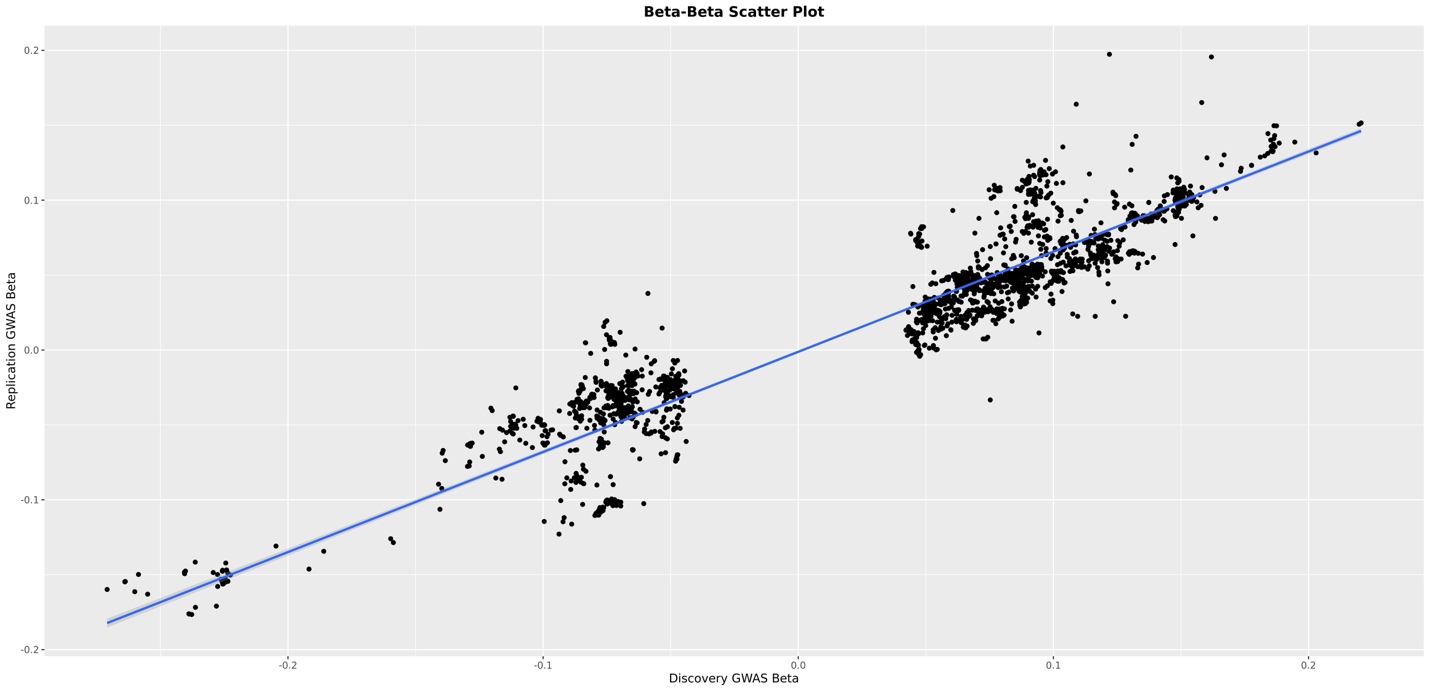
**Supplementary Figure 16:** A beta-beta plot showing the correlation between single nucleotide polymorphisms (SNPs) betas which were genome wide significant in our discovery GWAS, and their beta values in the replication GWAS.

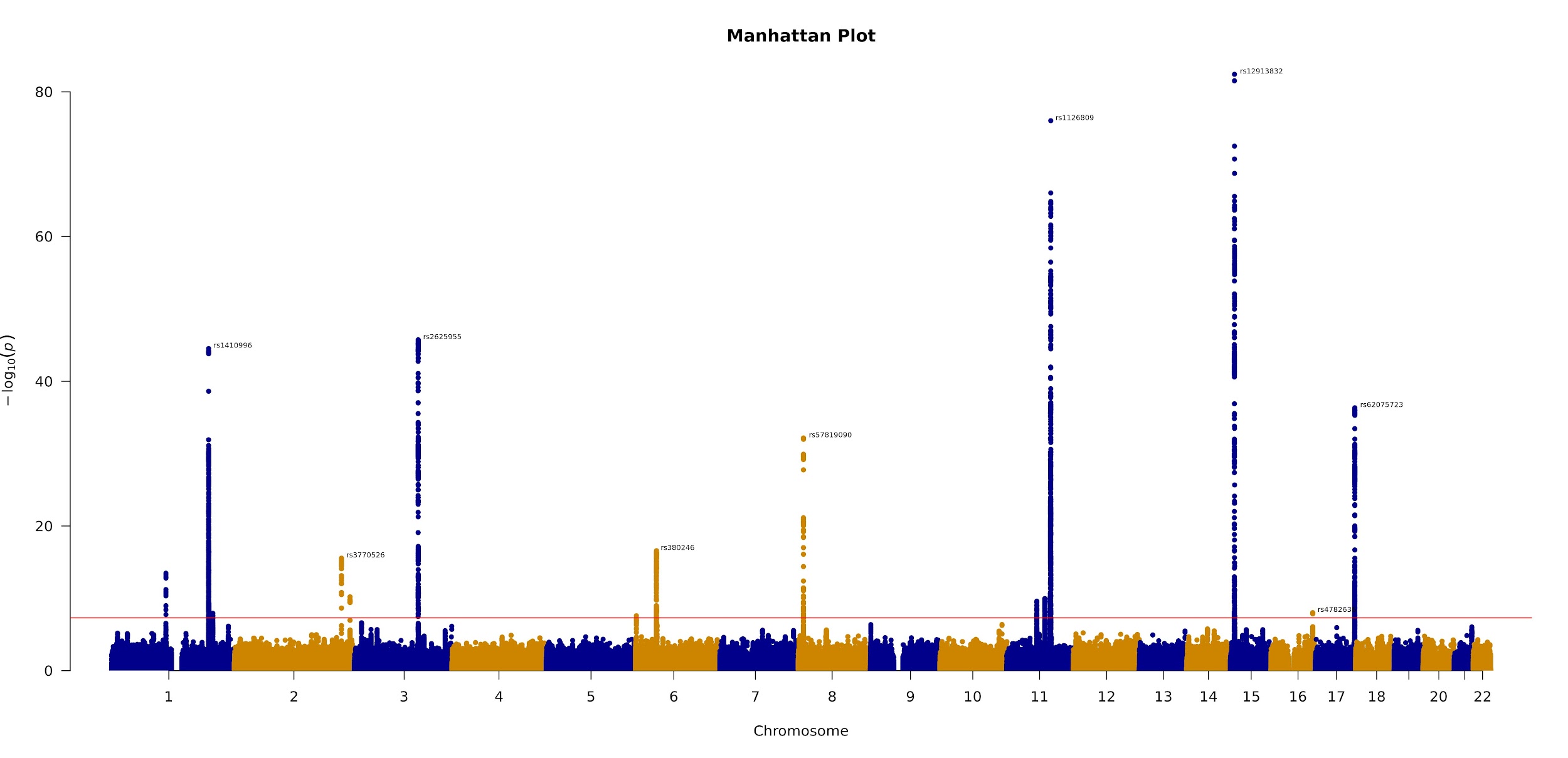
**Supplementary Figure 17**: The Manhattan plot for our GWAS of left eye overall RPE thickness. Genome-wide significant SNPs are annotated.
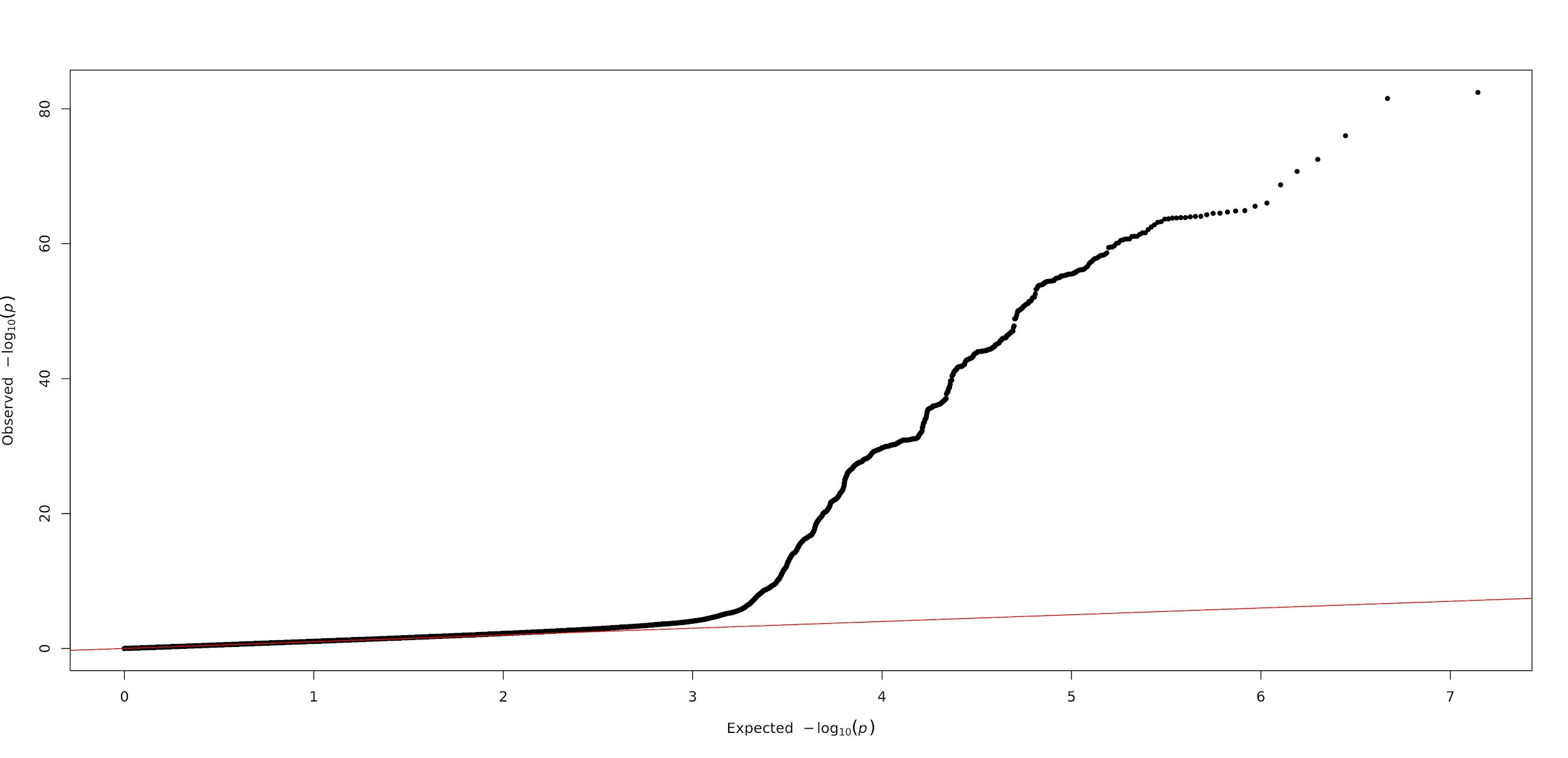

**Supplementary Figure 18**: The QQ plot for our GWAS of left eye overall RPE thickness.

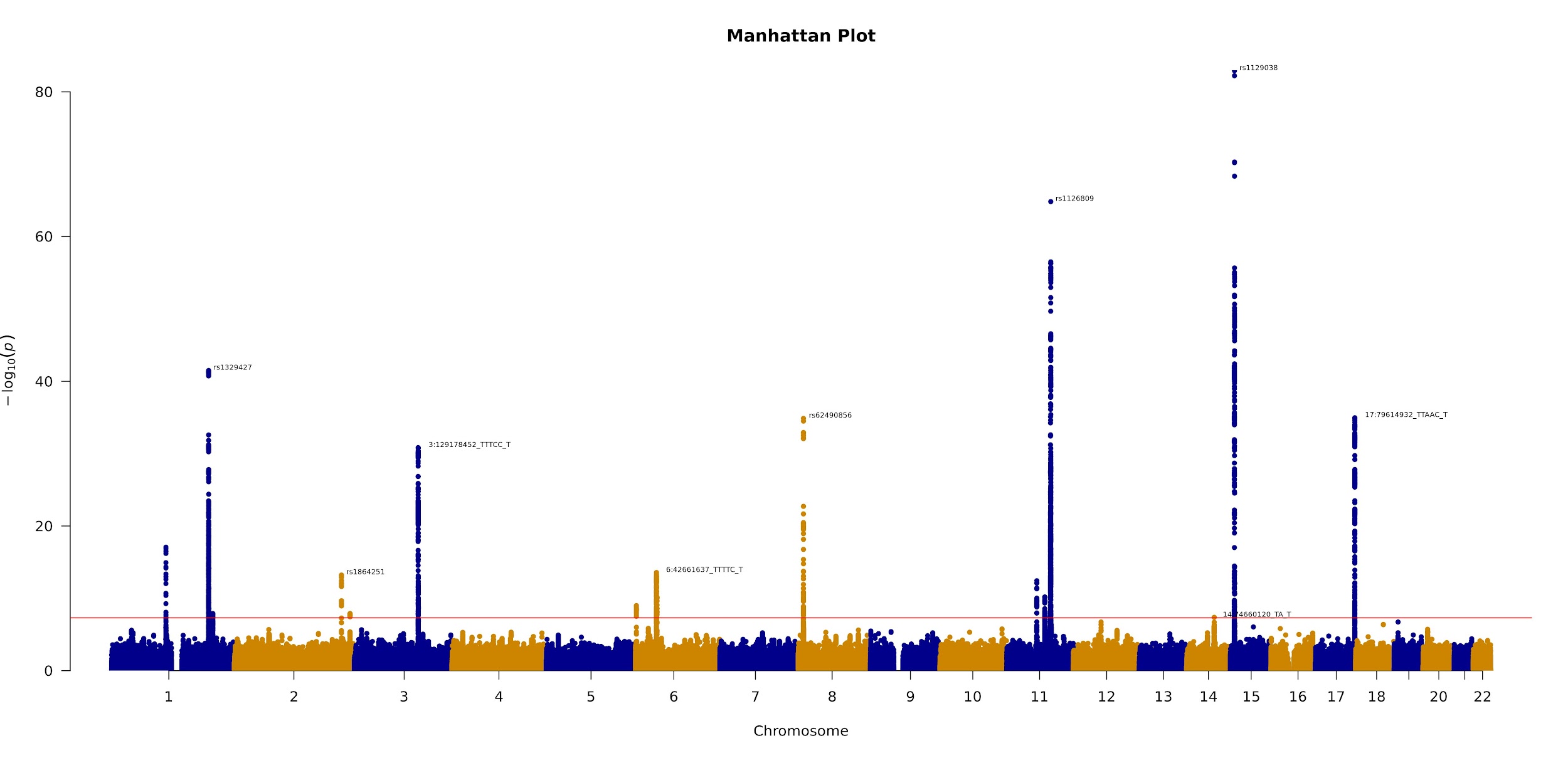
**Supplementary Figure 19**: The Manhattan plot for our GWAS of right eye overall RPE thickness. Genome-wide significant SNPs are annotated.
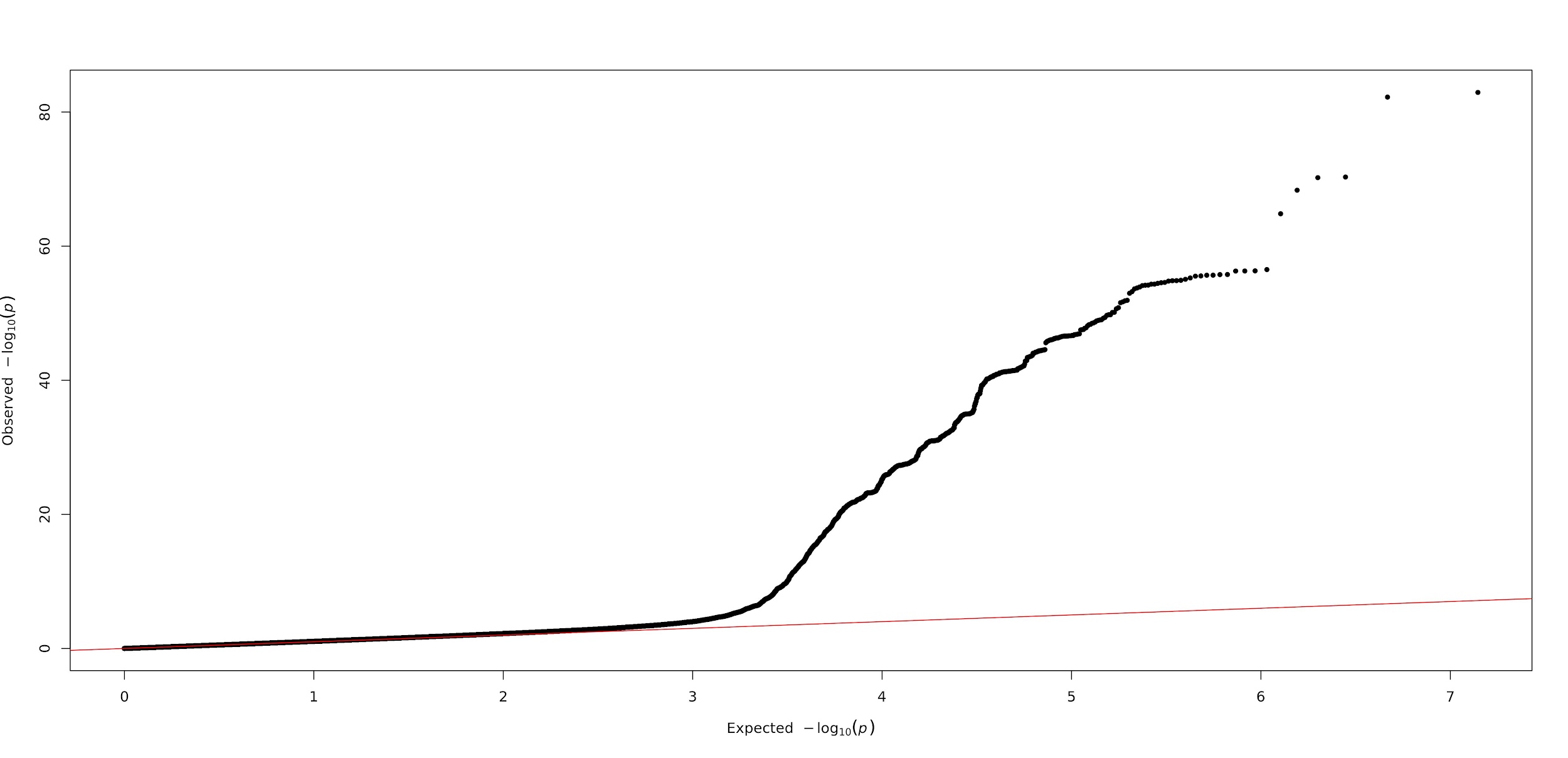

**Supplementary Figure 20**: The QQ plot for our GWAS of right eye overall RPE thickness.

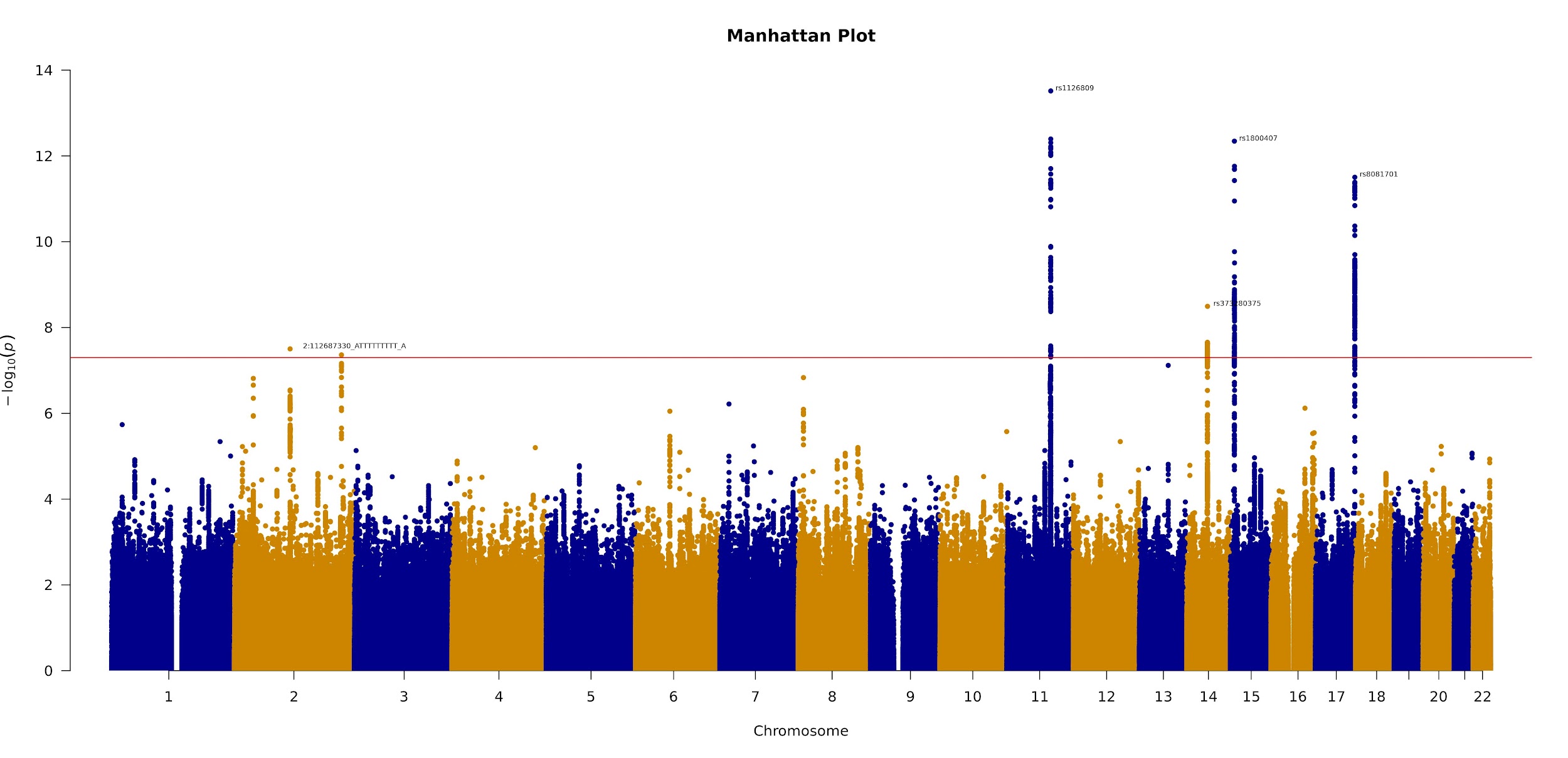

**Supplementary Figure 21:**  The Manhattan plot for our GWAS of left eye centre RPE thickness. Genome-wide significant SNPs are annotated.
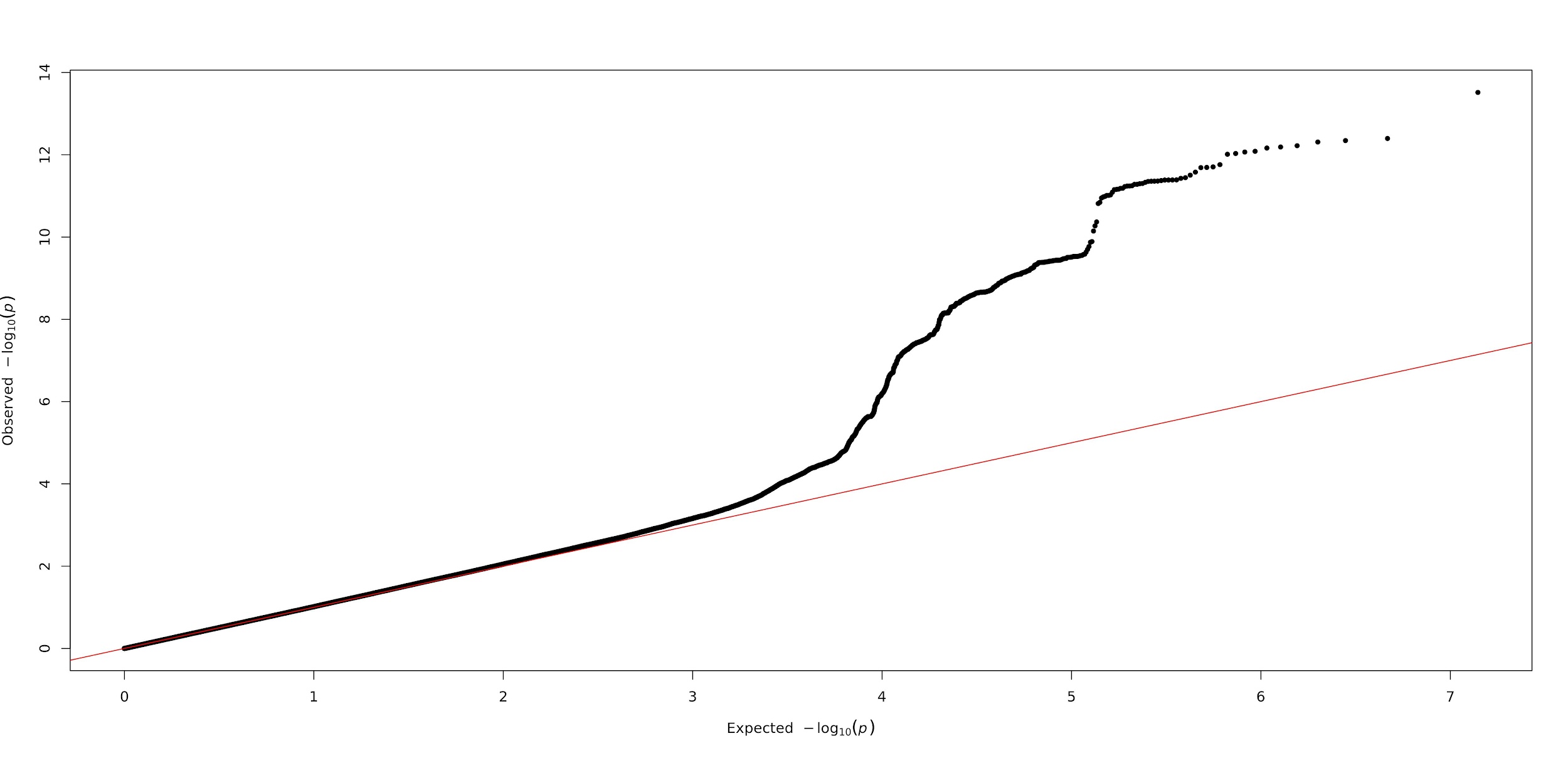

**Supplementary Figure 22**: The QQ plot for our GWAS of left eye centre RPE thickness.

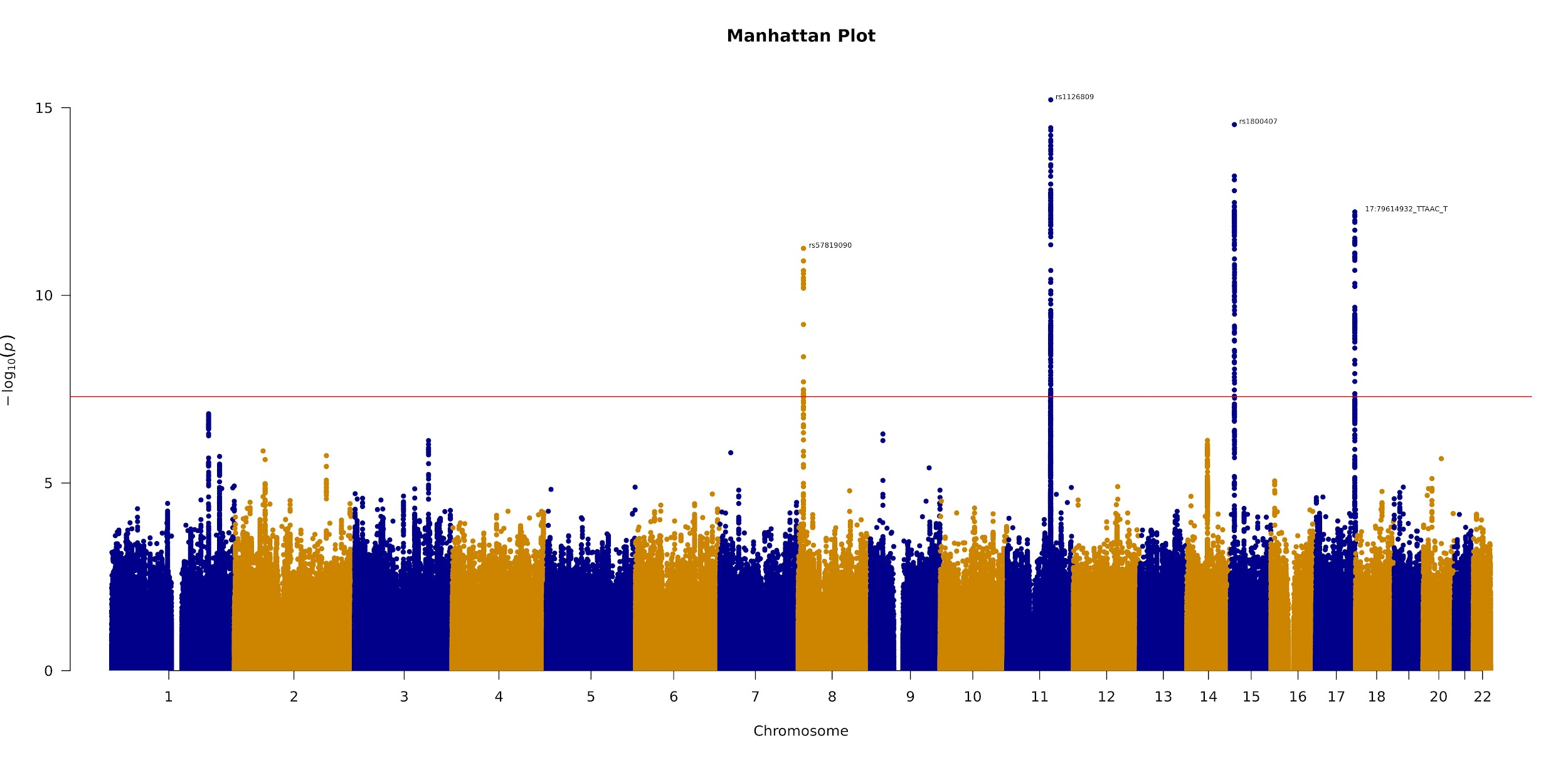
**Supplementary Figure 23:** The Manhattan plot for our GWAS of right eye centre RPE thickness. Genome-wide significant SNPs are annotated.
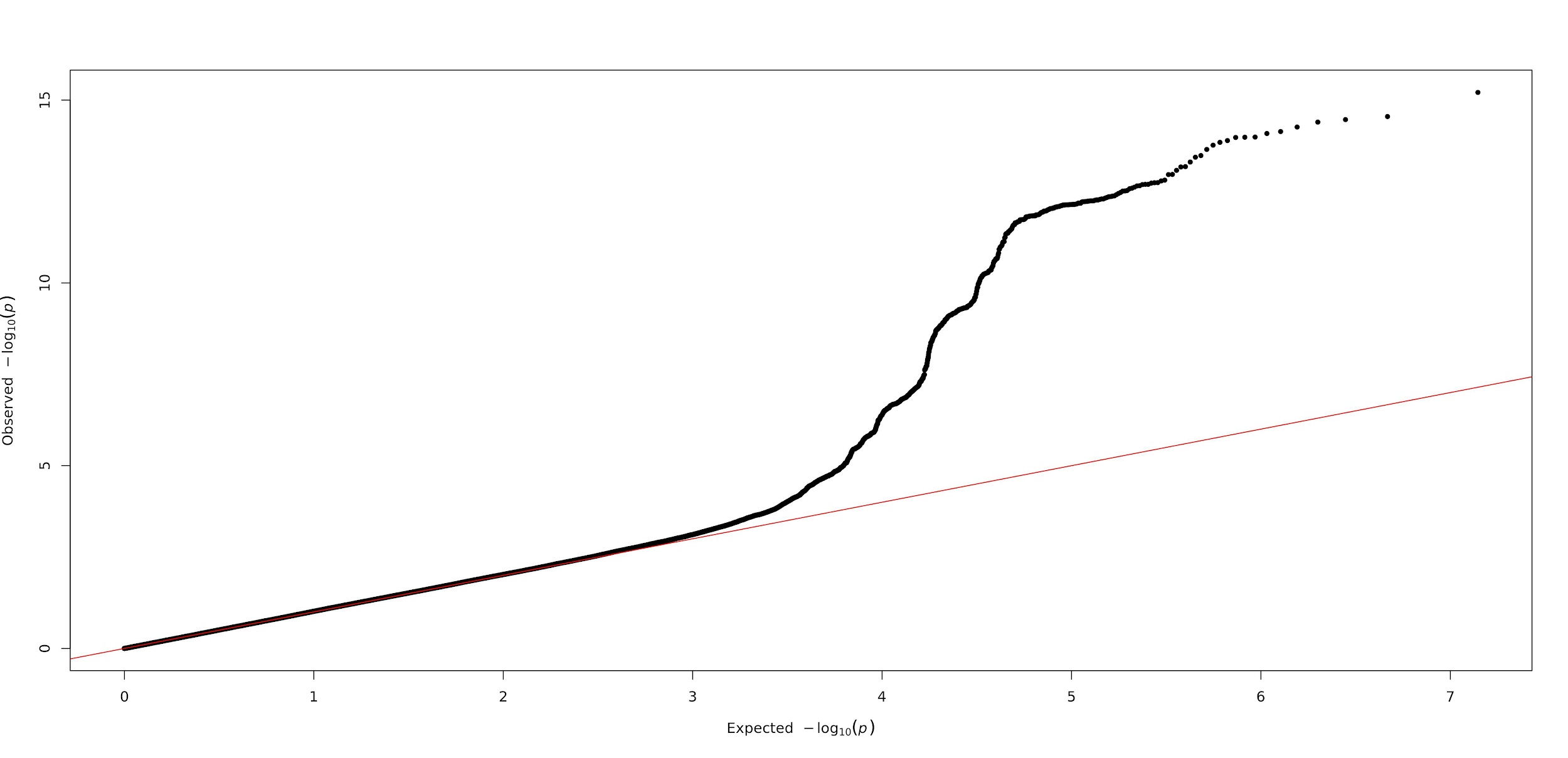
**Supplementary Figure 24:**  The QQ plot for our GWAS of right eye centre RPE thickness.

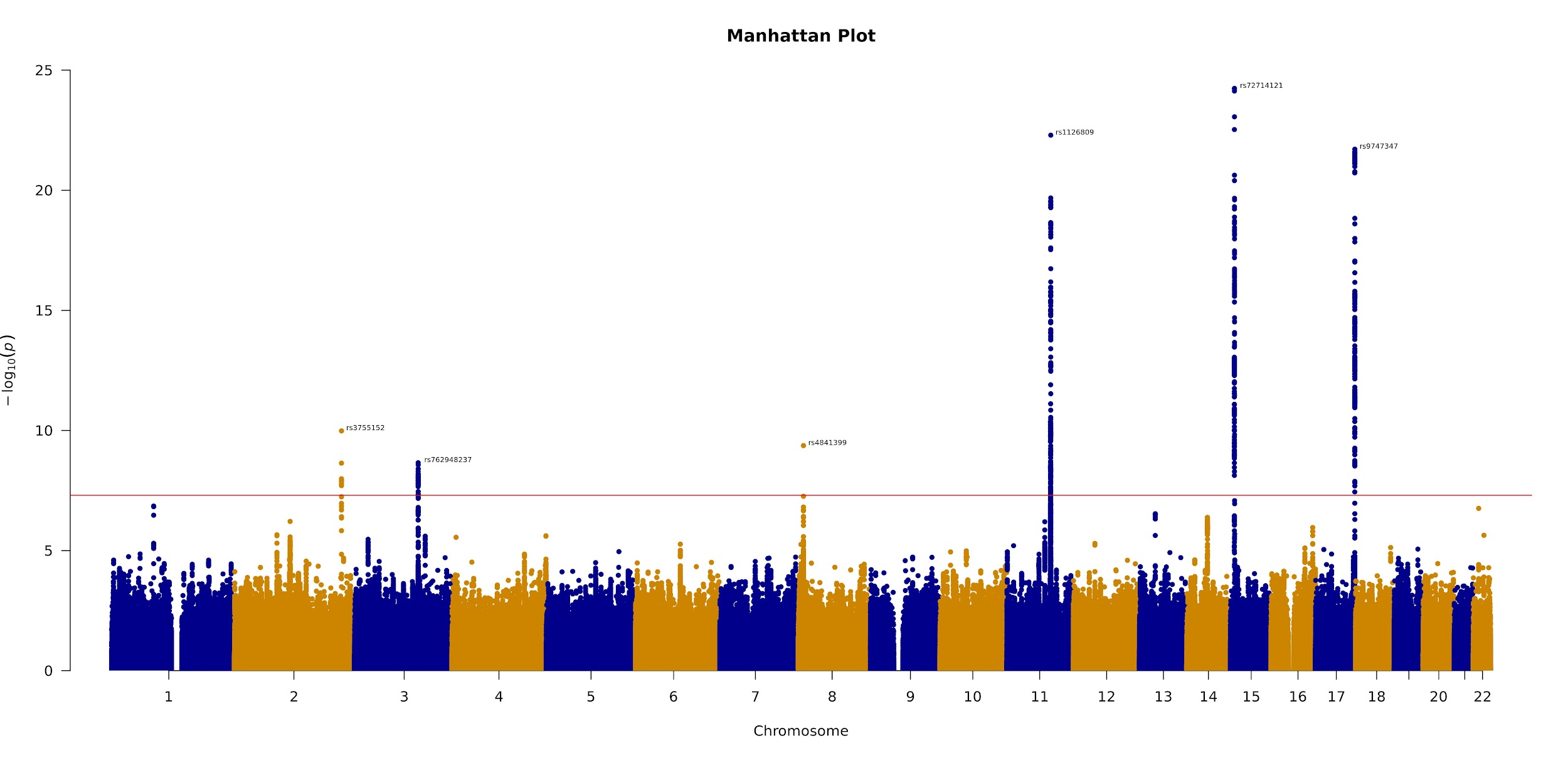

**Supplementary Figure 25:** The Manhattan plot for our GWAS of left eye mean inner RPE thickness. Genome-wide significant SNPs are annotated.

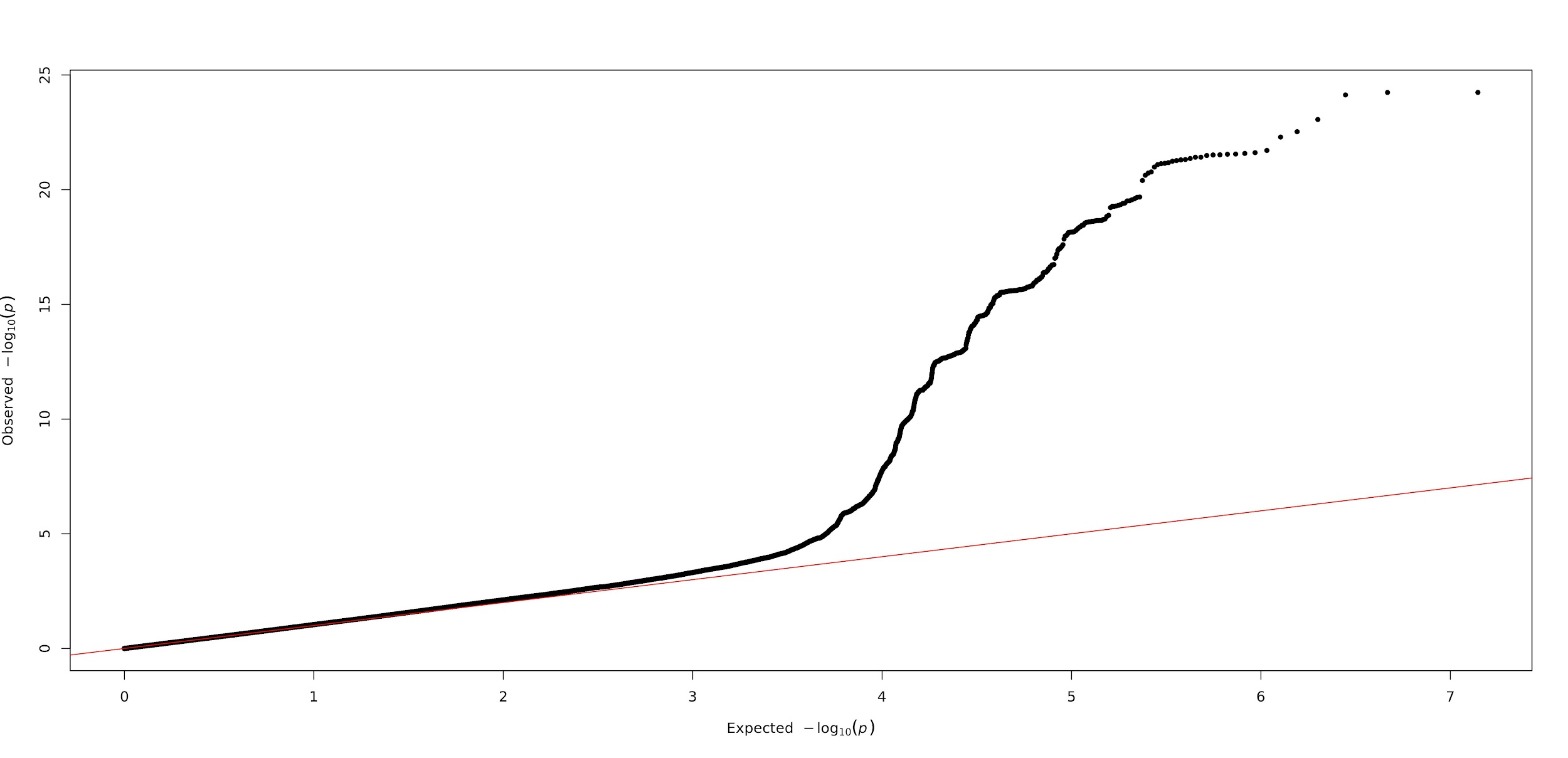

**Supplementary Figure 26:** The QQ plot for our GWAS of left eye mean inner RPE thickness.

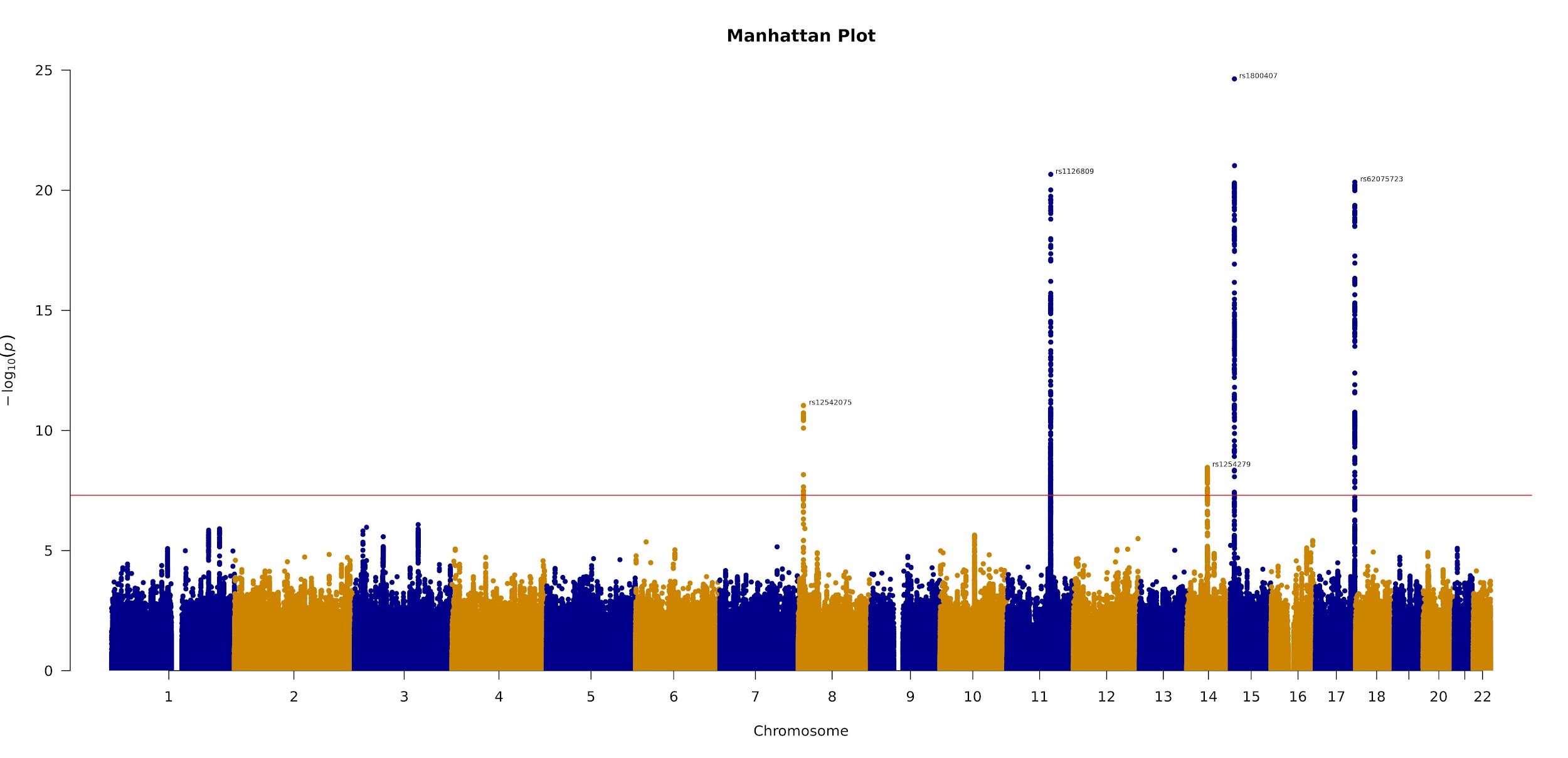

**Supplementary Figure 27:** The Manhattan plot for our GWAS of right eye mean inner RPE thickness. Genome-wide significant SNPs are annotated.

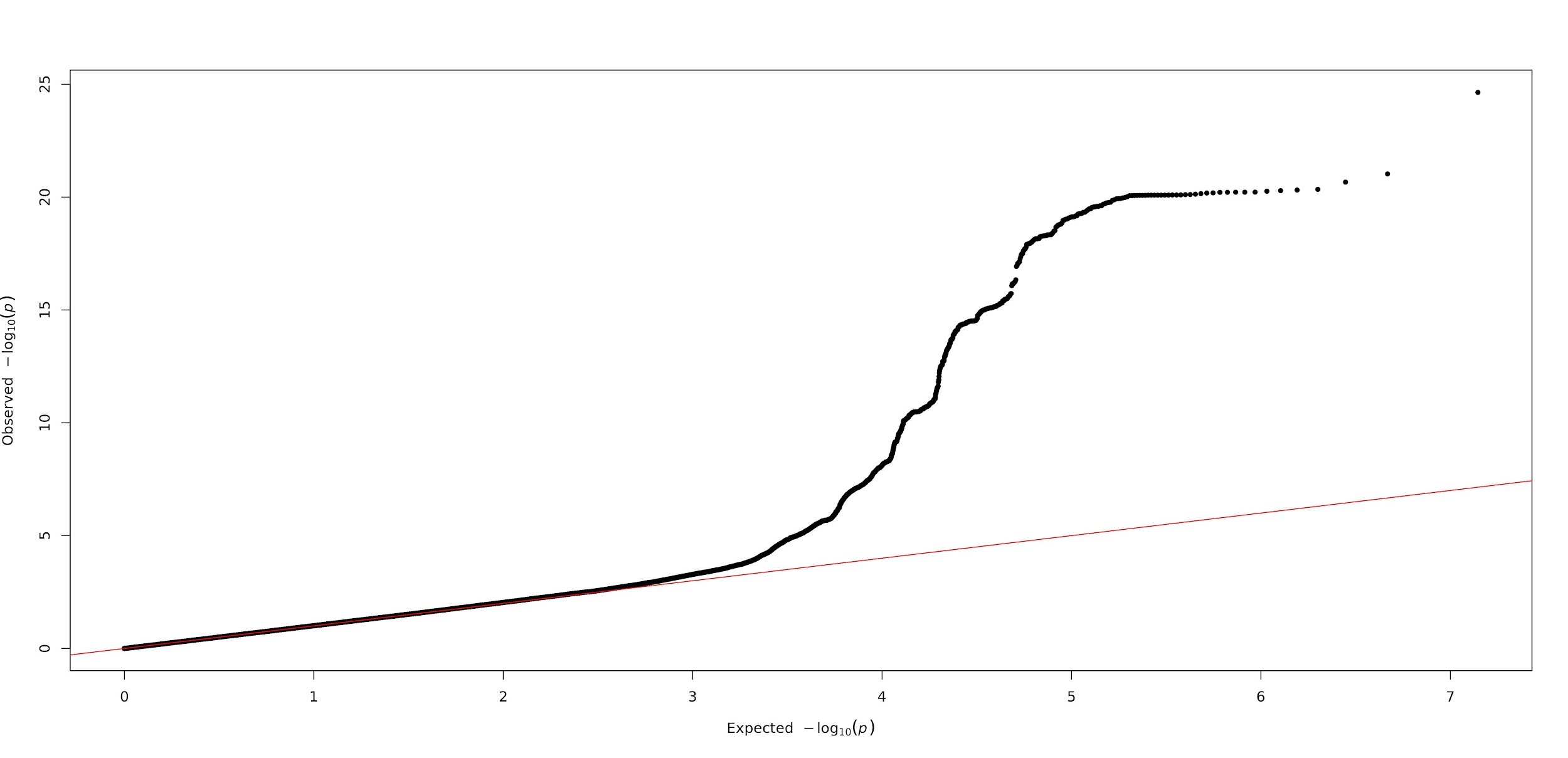

**Supplementary Figure 28:** The QQ plot for our GWAS of right eye mean inner RPE thickness.

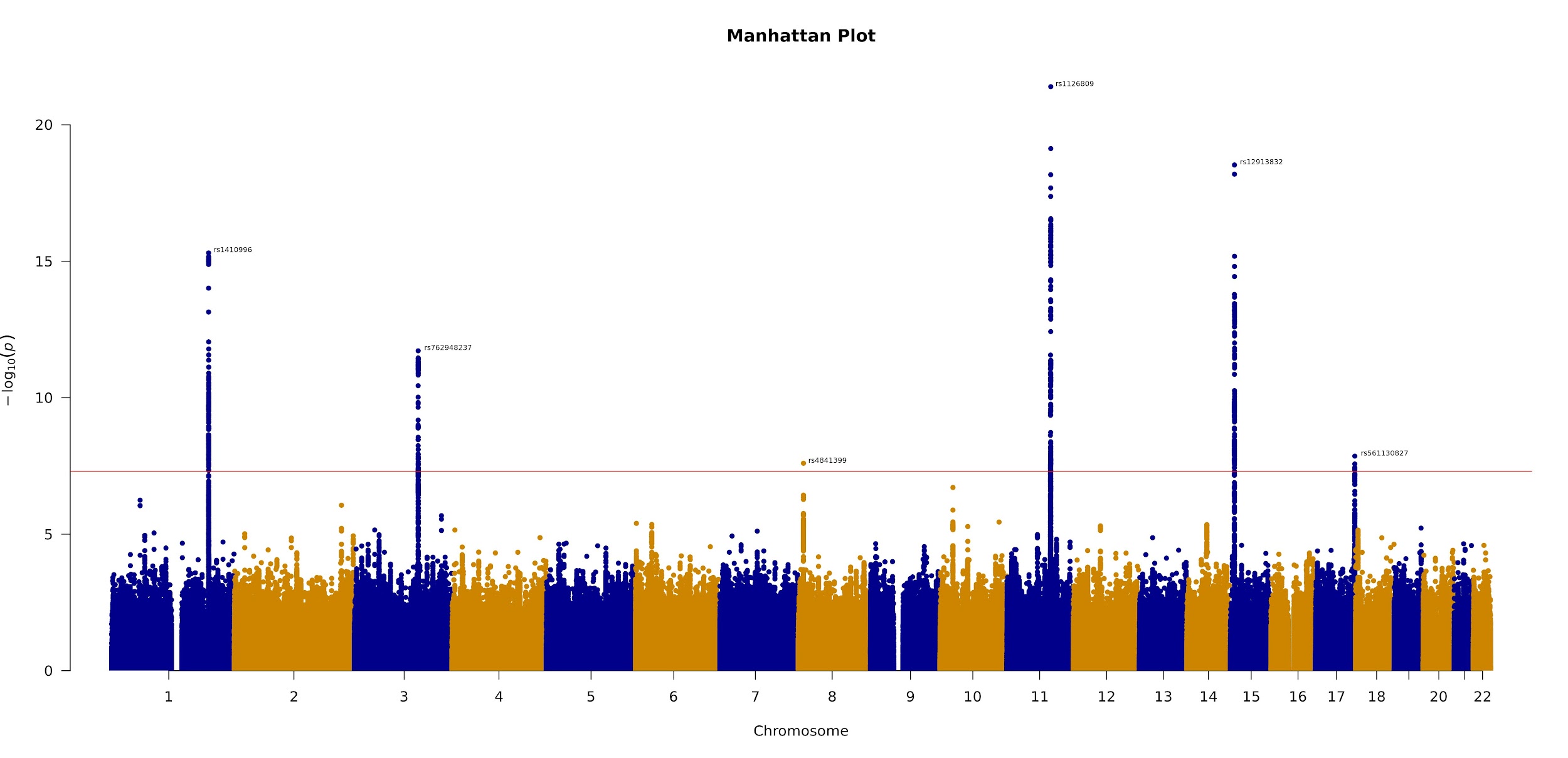

**Supplementary Figure 29:** The Manhattan plot for our GWAS of left eye mean outer RPE thickness. Genome-wide significant SNPs are annotated.

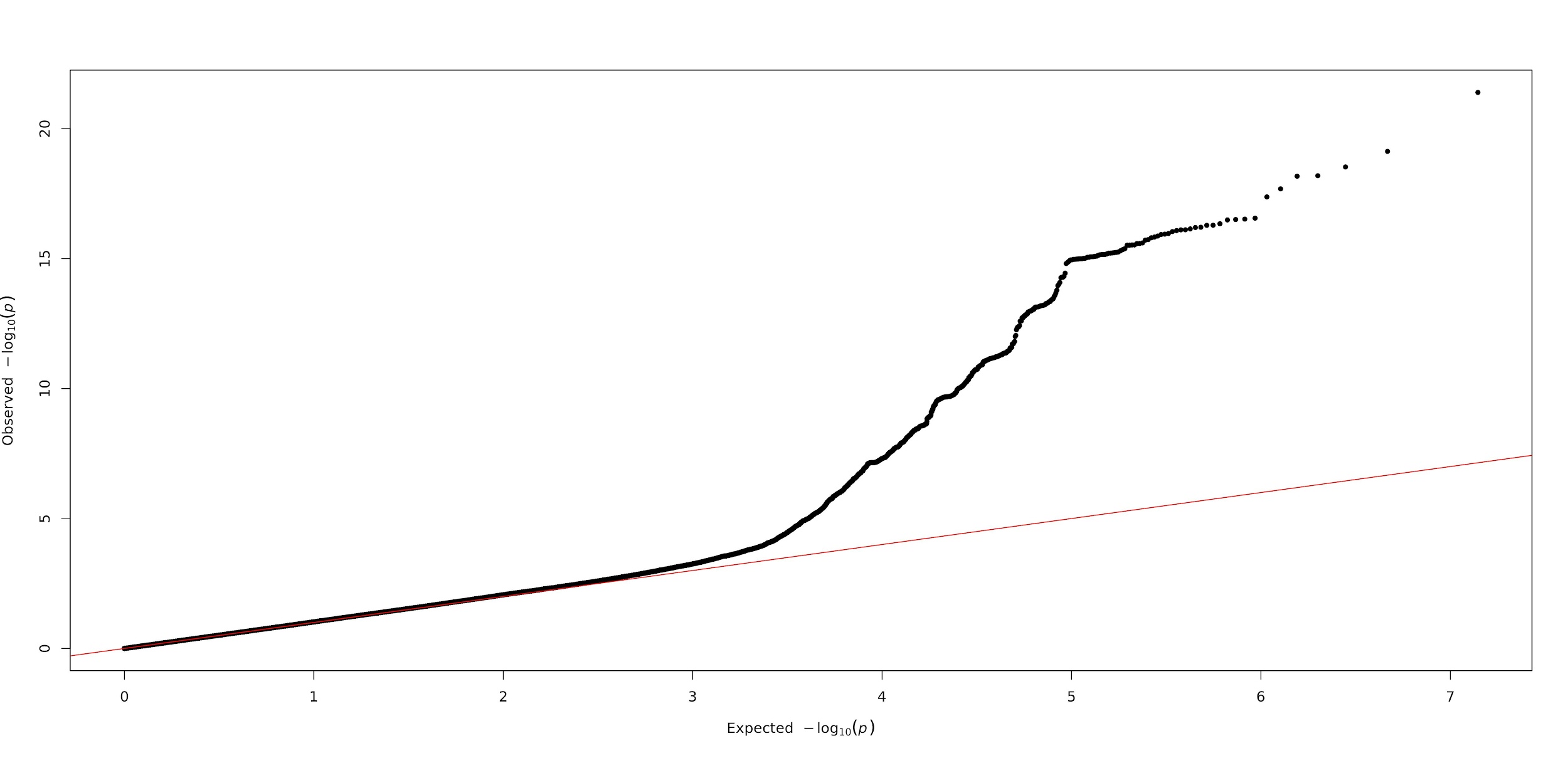

**Supplementary Figure 30:** The QQ plot for our GWAS of left eye mean outer RPE thickness.

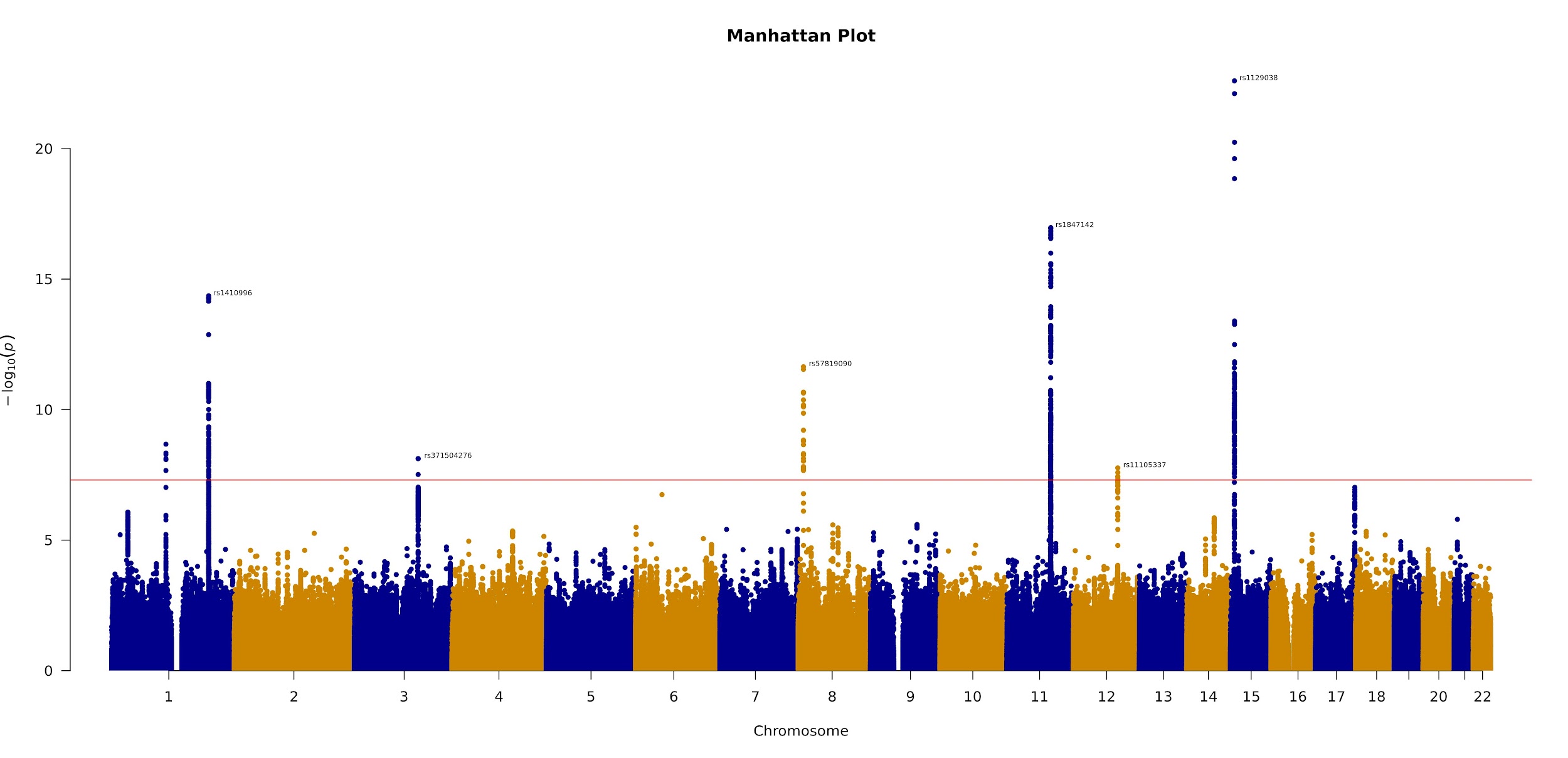

**Supplementary Figure 31:** The Manhattan plot for our GWAS of right eye mean outer RPE thickness. Genome-wide significant SNPs are annotated.

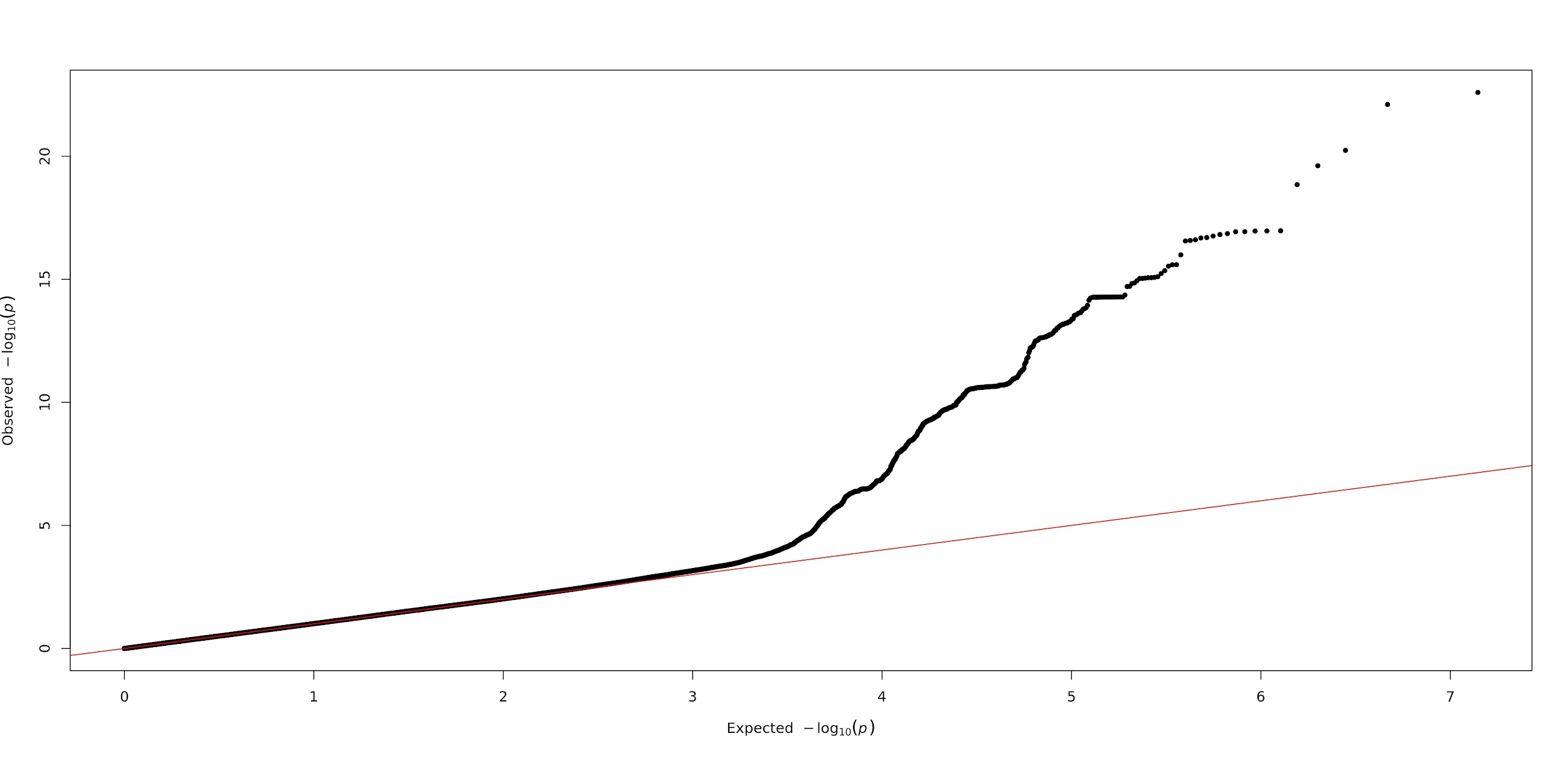

**Supplementary Figure 32:** The QQ plot for our GWAS of right eye mean outer RPE thickness.

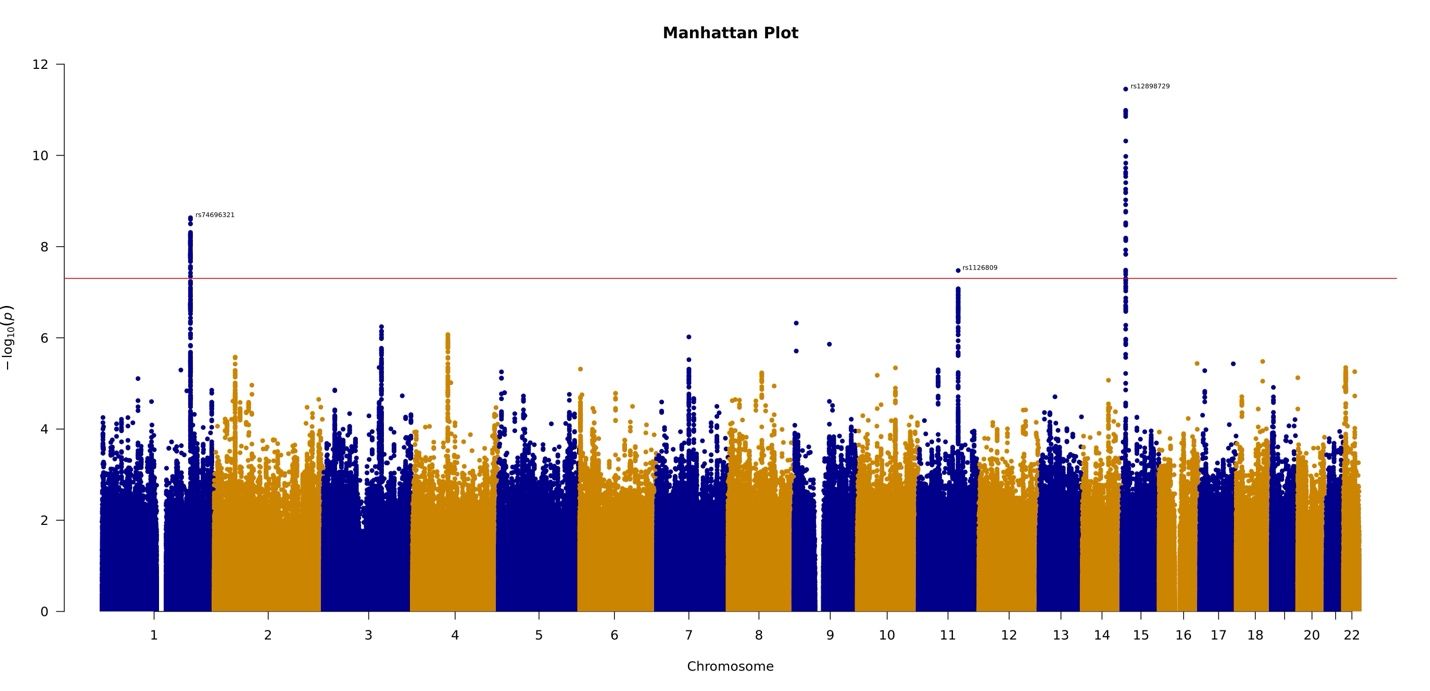

**Supplementary Figure 33:** The Manhattan plot for our GWAS of replication study of left eye overall RPE thickness.

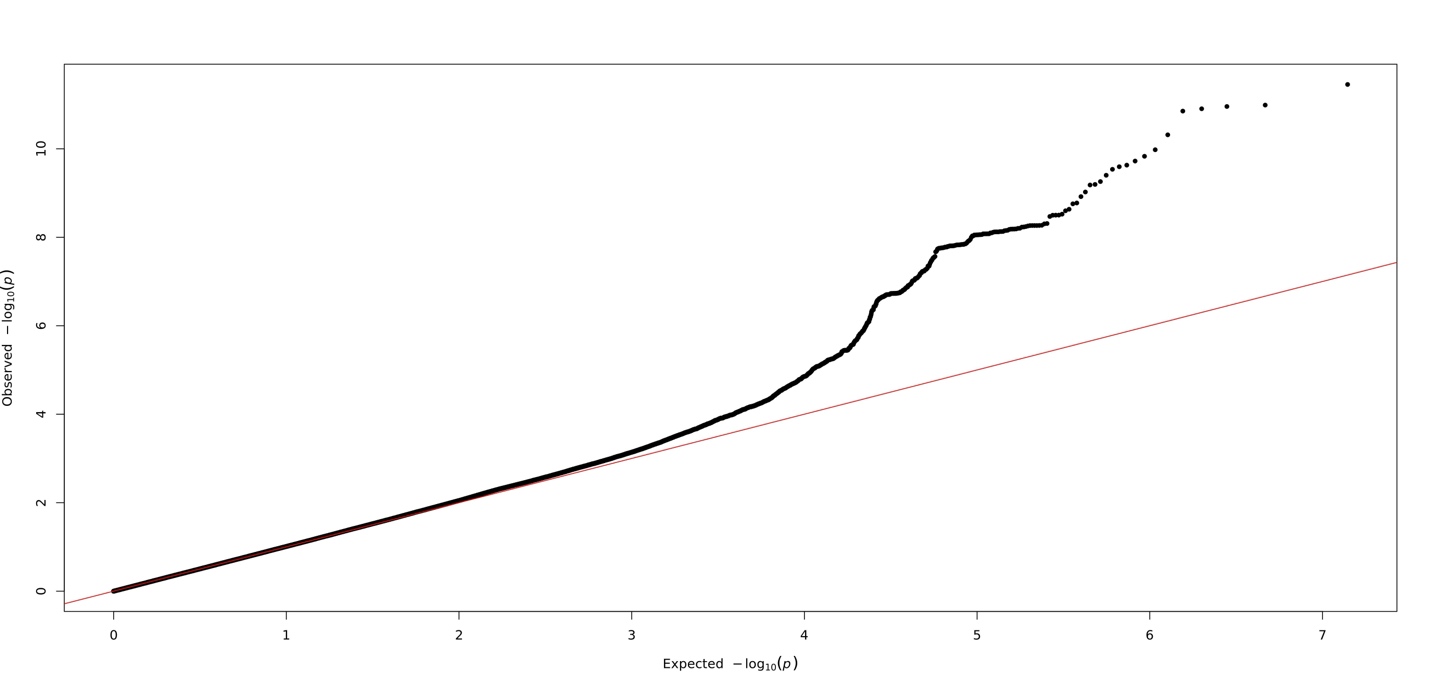

**Supplementary Figure 34:** The QQ plot for our GWAS of replication study of left eye overall RPE thickness.

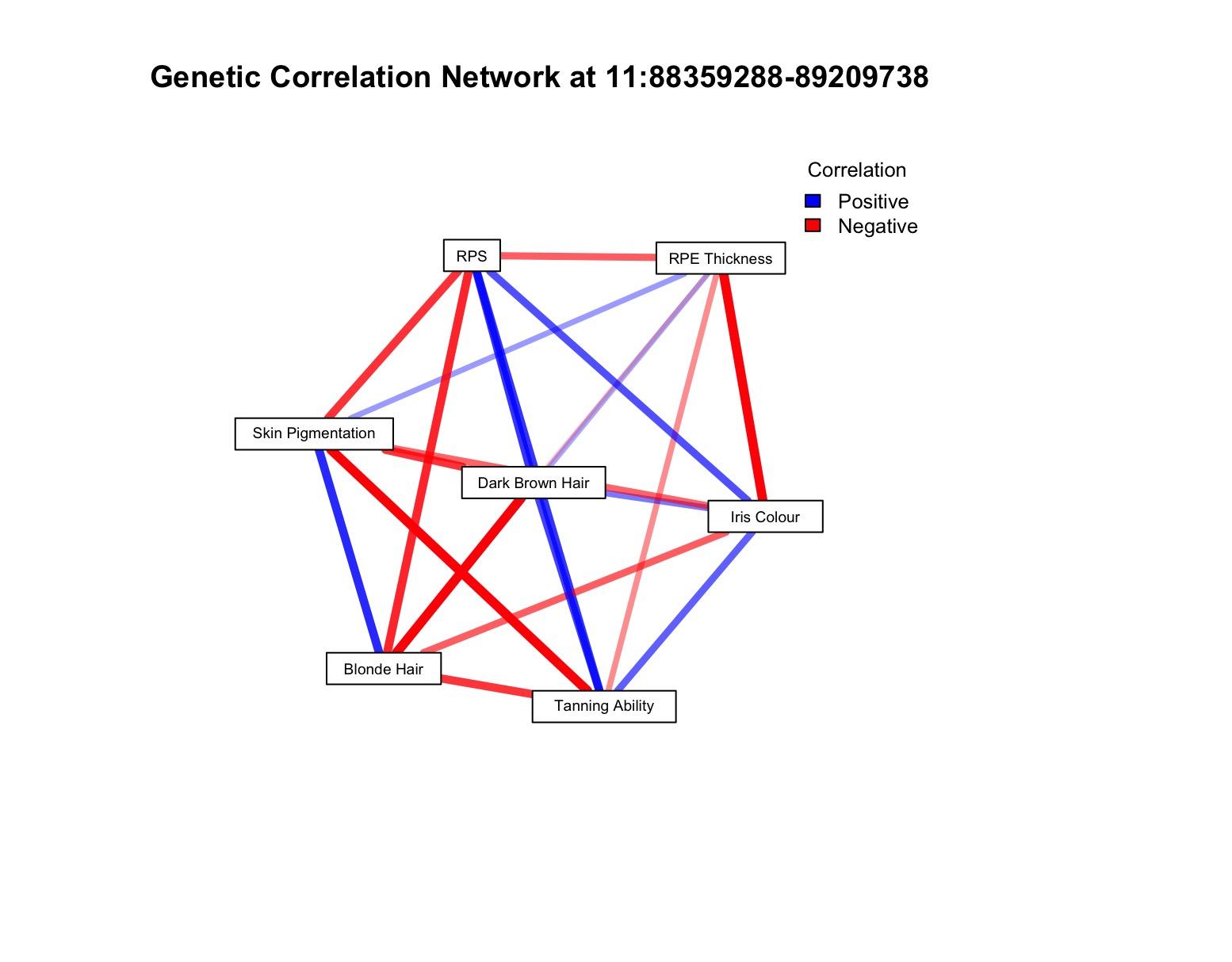

**Supplementary Figure 35:** Network chart detailing the local genetic correlations at 11:88359288-89209738, the locus containing *GABRG3, OCA2-HERC2, LOC645202, HERC2P11* and *GOLGA8M*. Blue connections indicate positive correlations whilst red indicates negative correlations. The opacity of the connection is greatest for weaker correlations in terms of coefficient, with stronger correlations indicated by wider, solid lines. All connections represent multiple testing corrected significant local genetic correlations.

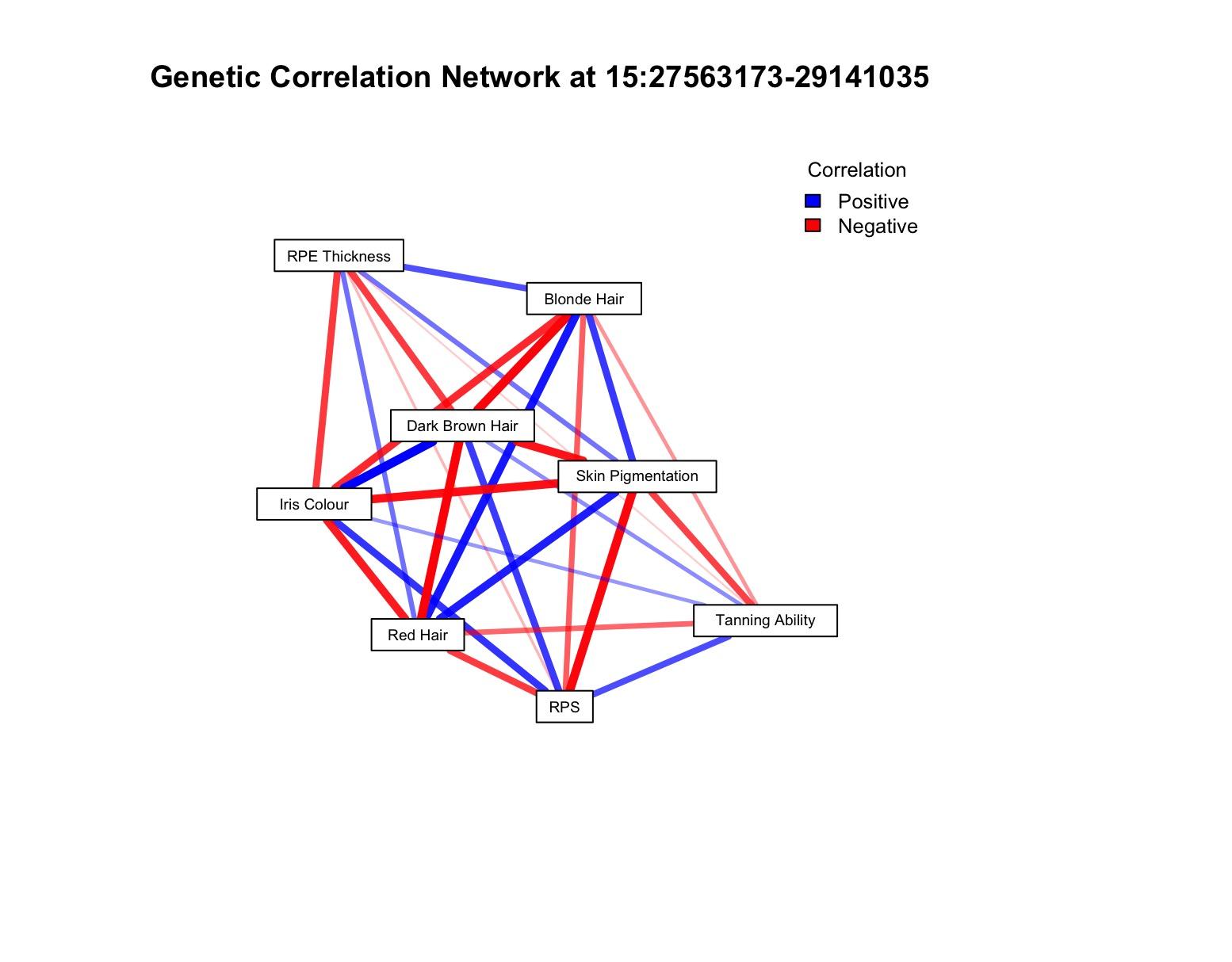

**Supplementary Figure 36:** Network chart detailing the local genetic correlations at 11:88359288-89209738, the locus containing *GRM5, TYR* and *NOX4*. Blue connections indicate positive correlations whilst red indicates negative correlations. The opacity of the connection is greatest for weaker correlations in terms of coefficient, with stronger correlations indicated by wider, solid lines. All connections represent multiple testing corrected significant local genetic correlations.
